# Acute GARP depletion disrupts vesicle transport, leading to severe defects in sorting, secretion, and O-glycosylation

**DOI:** 10.1101/2024.10.07.617053

**Authors:** Amrita Khakurel, Irina Pokrovskaya, Vladimir V. Lupashin

## Abstract

The GARP complex is an evolutionarily conserved protein complex proposed to tether endosome-derived vesicles at the trans-Golgi network. While prolonged depletion of GARP leads to severe trafficking and glycosylation defects, the primary defects linked to GARP dysfunction remain unclear. In this study, we utilized the mAID degron strategy to achieve rapid degradation of VPS54 in human cells, acutely disrupting GARP function. This resulted in the partial mislocalization and degradation of a subset of Golgi-resident proteins, including TGN46, ATP7A, TMEM87A, CPD, C1GALT1, and GS15. Enzyme recycling defects led to the early onset of O-glycosylation abnormalities. Additionally, while the secretion of fibronectin and cathepsin D was altered, mannose-6-phosphate receptors were largely unaffected. Partial displacement of COPI, AP1, and GGA coats caused a significant accumulation of vesicle-like structures and large vacuoles. Electron microscopy detection of GARP-dependent vesicles, along with the identification of specific cargo proteins, provides direct experimental evidence of GARP’s role as a vesicular tether. We conclude that the primary defects of GARP dysfunction involve vesicular coat mislocalization, accumulation of GARP-dependent vesicles, degradation and mislocalization of specific Golgi proteins, and O-glycosylation defects.

## Introduction

Proteins and lipids within the cell are continuously trafficked between the plasma membrane and the *trans*-Golgi network (TGN) via the endosome-to-TGN pathway [1] [2]. This retrograde transport mechanism is crucial for the recycling of protein and lipid cargoes, balancing the anterograde movement of membranes [3] and preventing the degradation of these components in lysosomes [4]. Some of the cargoes that utilize endosome-to-TGN trafficking include the copper transporters ATP7A and ATP7B [5] [6] [7], enzymes carboxypeptidase D and furin [8] [9] [10], putative ion channel TMEM87 [3] and recycling receptors such as mannose-6-phosphate receptors (MPRs) [11], sortilins [12] [13] and TGN46 [1]. This trafficking step is also exploited by multiple pathogens, including cholera [14], Shiga [15] and SubAB [16] toxins.

Cargo transport between cellular compartments begins with the selection and packaging of cargo into small membrane intermediates (vesicles or tubules) at the donor compartment [17], and ends with the tethering of these vesicles and their subsequent fusion with the acceptor compartment [18] [19]. At the TGN, cargo-laden vesicle tethering is mediated by long coiled-coil proteins known as Golgins [20] [21] and the multisubunit tethering complex (MTC) Golgi Associated Retrograde Protein (GARP) [22] [11] [23]. The GARP complex is evolutionarily conserved across a range of organisms, including humans, mice, and plants [24] [25] [26]. GARP belongs to the CATCHR (Complexes associated with tethering containing helical rods) family of MTCs and is thought to tether retrograde transport vesicles originated from endosomes, facilitating their fusion with the TGN [27] [28] [29]. The GARP complex is composed of four subunits: VPS51, VPS52, VPS53, and VPS54 [23]. Of these, VPS51, VPS52, and VPS53 are shared with the EARP (endosome-associated recycling protein) complex, while VPS54 is unique to GARP [30]. In mammalian cells, GARP’s localization to the TGN relies on small GTPases ARFRP1 and ARL5 [31]. GARP role in retrograde trafficking is supported by multiple interactions with other components of the endosome-TGN trafficking machinery [32] [33] [34]. However, the mechanism of GARP’s action remains unclear. Mutations in the VPS51 have been associated with abnormal glycosylation patterns in patients [30]. Similarly, knockout (KO) of VPS53 and VPS54 in tissue culture cells causes severe defects in both N- and O-linked protein glycosylation resulting from mislocalisation and degradation of multiple Golgi enzymes [35] [36]. Moreover, GARP-KO led to significant mislocalization of COPI, AP1, and GGA vesicle coats, displacement of ARF1 GEFs (GBF1 and BIG1), and severe alterations in Golgi morphology. Although the expression of missing GARP subunits rescues all observed defects, some of these defects may be secondary, arising from the persistent mistargeting of receptors and cellular trafficking machinery, or cellular adaptation to the chronic loss of the GARP complex.

To investigate the primary defects caused by GARP dysfunction, we developed a novel cellular system that enables the acute depletion of VPS54, a key subunit of the GARP complex, using the auxin-inducible degron (mAID) technology [37] [38] [39] [40]. A combination of biochemical and microscopic techniques was used to analyze the impact of acute VPS54 depletion on Golgi morphology, stability of other GARP subunits, GARP-interacting membrane trafficking partners, glycosylation enzymes, and other Golgi resident proteins. This study provides a comprehensive view of the primary cellular defects associated with GARP dysfunction in human cells.

## Materials and Methods

### Cell Culture

hTERT-RPE1 (retinal pigment epithelial, RPE1) and HEK293T cells used for all experiments were purchased from ATCC. RPE1 VPS54-KO cells were described previously [36]. HeLa-KO cells were obtained from Bonifacino lab (NIH) [31]. RPE1, HEK293T and HeLa cells were cultured in Dulbecco’s Modified Eagle’s Medium (DMEM) containing Nutrient mixture F-12 (Corning) supplemented with 10% fetal bovine serum (FBS) (Thermo Fisher). Cells were incubated in a 37°C incubator with 5% CO2 and 90% humidity. All DNA plasmids used in this work are listed in Tables 1.

**Table 1.**
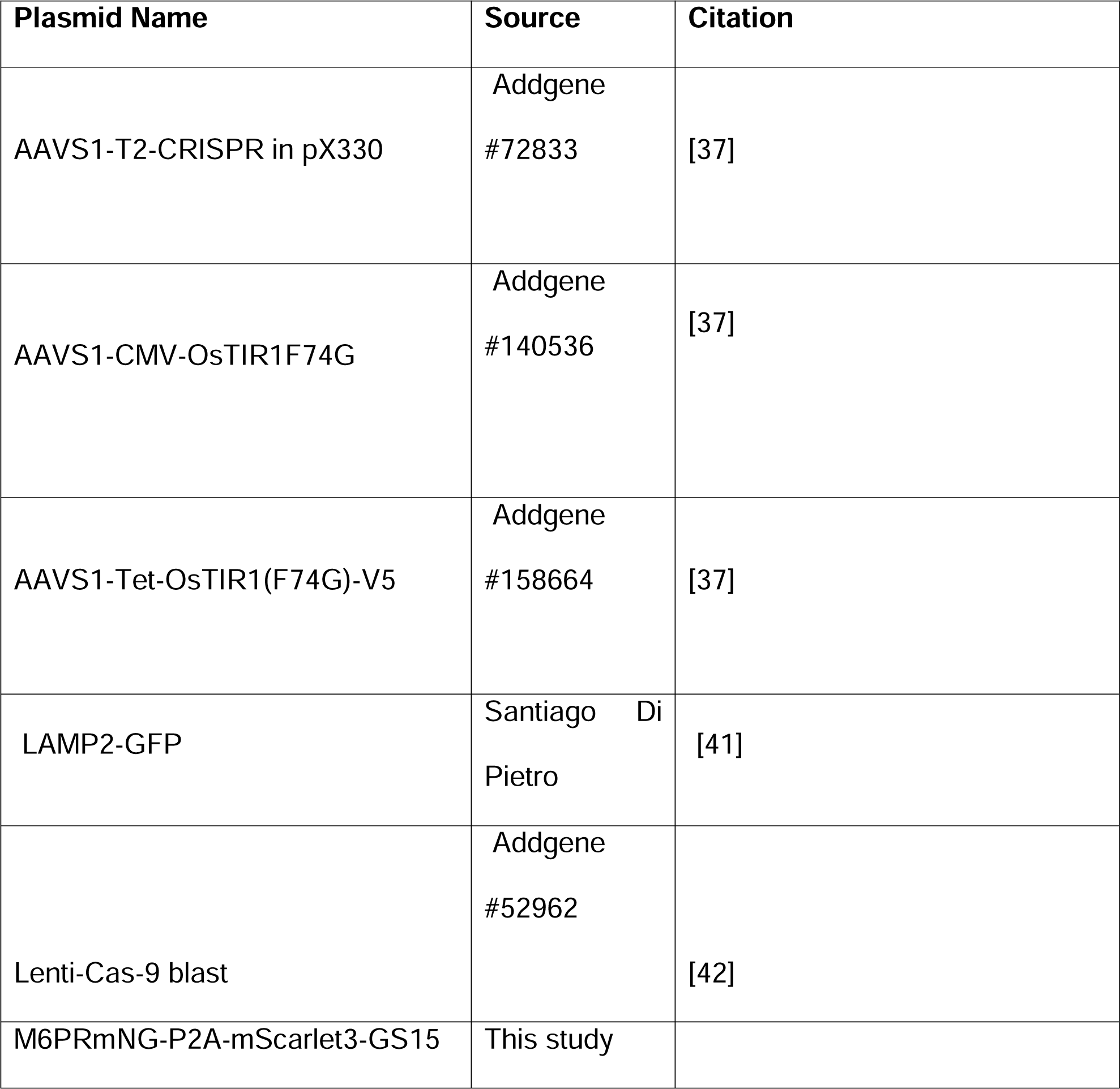
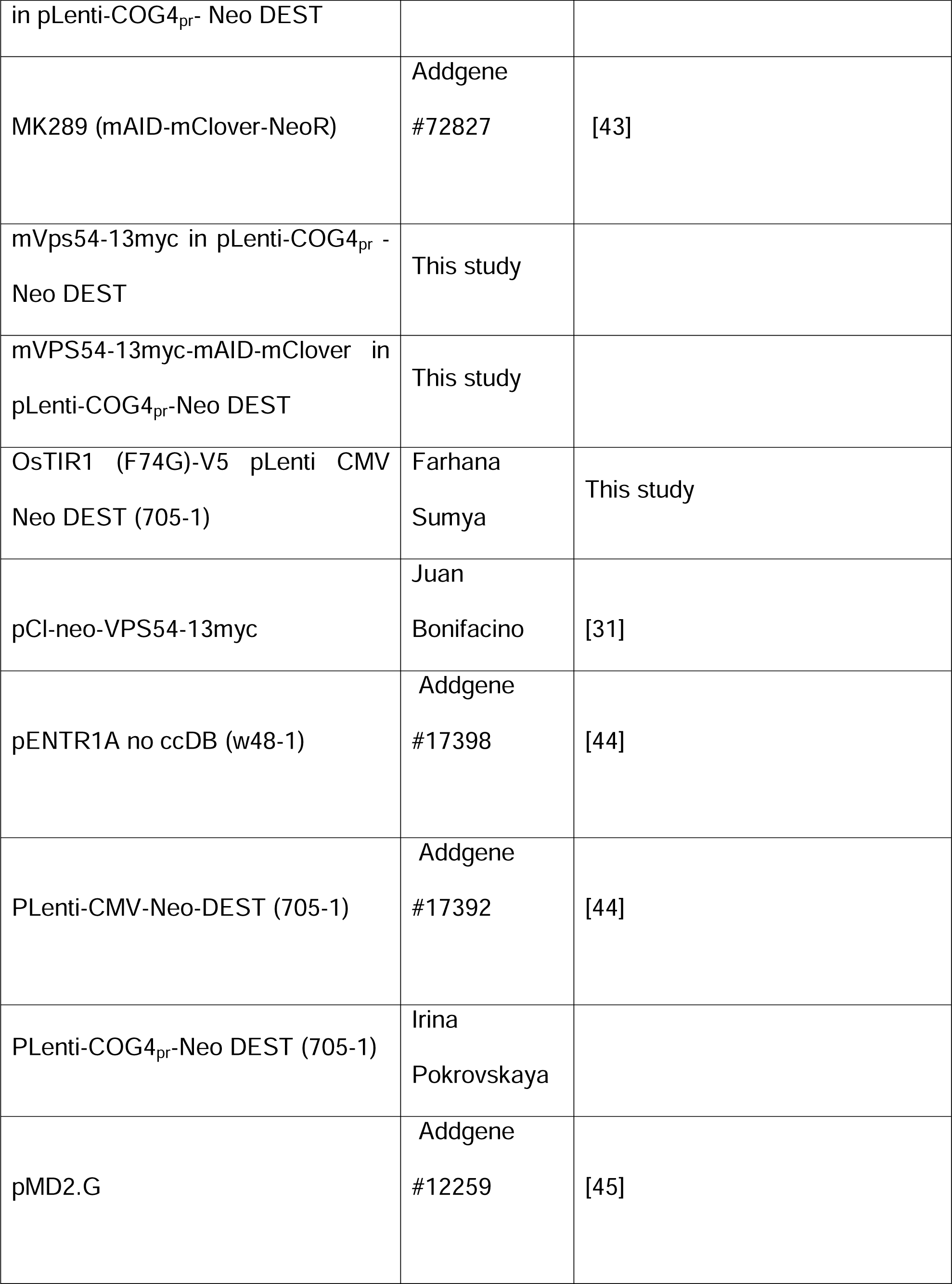
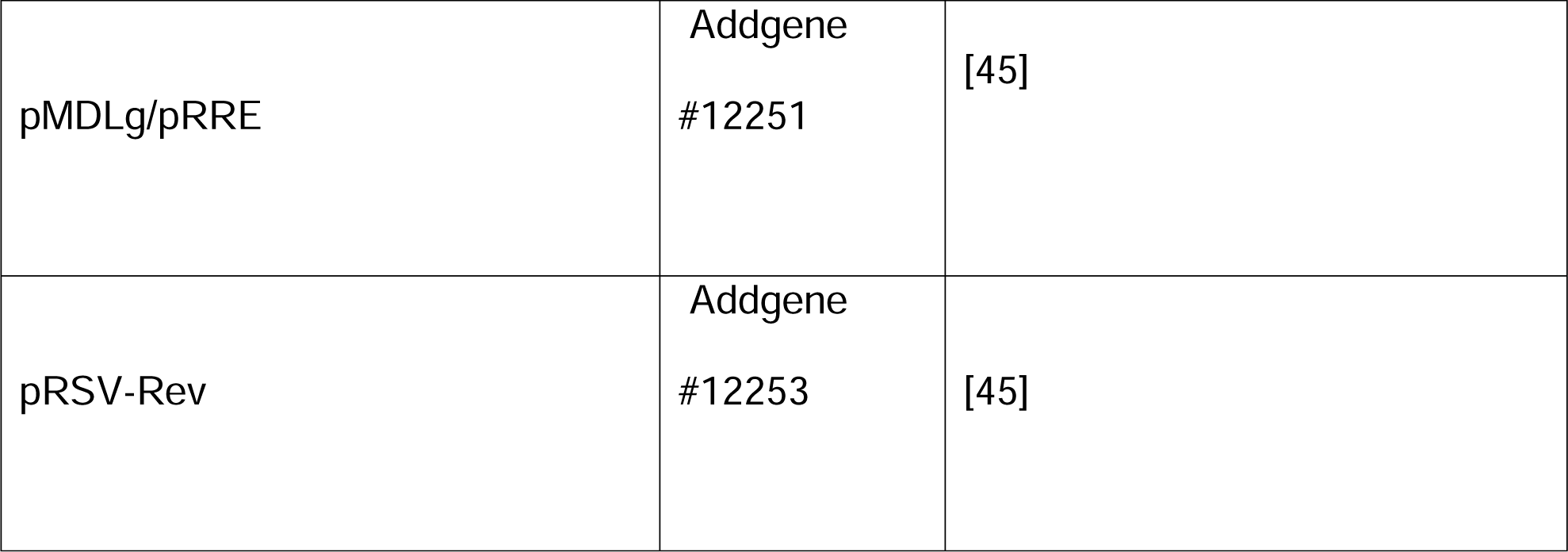
List of plasmids used in the study.

### Preparation of mVPS54-mAID expressing cells

hTERT RPE1 VPS54-KO cells were rescued with mVPS54-13-myc–mAID-mClover. For convenience, we will use VPS54-mAID hereafter.

Briefly, mVPS54 in pENTR1A 48-1 was amplified using VPS54-Xba1-Forward (CGGCCGCACTCGAGATATCTAGACCCAG) and VPS54-BamH1-Reverse (ATTGGATCCGTGGTGATGGTGGTGGTGATG) primers.

PCR product was purified using the QIAquick PCR Purification Kit (QIAGEN) following the standard protocol. To create VPS54-mAID in pENTR1A, mVPS54 in pENTR1A (48-1) and MK289 (mAID-mClover-NeoR) were digested with BamHI and XbaI, and ligated. This construct was then recombined with the pLentiCOG4_pr_-Neo–DEST plasmid, using Gateway LR Clonase II Enzyme Mix (Thermo Fisher). The recombined plasmid was transformed into Stbl3 competent cells as per the manufacturer’s instructions, and DNA was extracted using the QIAprep Spin Miniprep Kit. VPS54-mAID pLenti clones were verified by restriction analysis. The expression of mVPS54-mAID was validated by transfecting HEK293T cells with the selected pLenti plasmids and performing Western blot (WB) analysis using an anti-myc antibody.

To produce lentiviral particles, HEK293FT cells were co-transfected with equal amounts of lentiviral packaging plasmids (pMD2.G, pRSV-Rev, pMDLg/pRRE) and the mVPS54-mAID pLenti plasmid using Lipofectamine 3000, following the manufacturer’s protocol as previously described [36]. hTERT-RPE1 VPS54-KO cells were transduced with the lentivirus expressing mVPS54-mAID. Single-cell clones were isolated by serial dilution, expanded, and validated by WB and immunofluorescence (IF) for stable expression of mVPS54-mAID.

### Construction of cells that co-express mVPS54-mAID and OsTIR1 (F74G)-V5

hTERT-RPE1 VPS54-KO cells expressing mVPS54-mAID were transduced with lentiviral AAVS1 CMV-OsTIR1F74G. Briefly, OsTIR1 (F74G)-V5 was amplified using OsTIR1 (F74G)-V5 SAL1 Forward (GAGGTCGACATGACATACTTTCCTGAAGA) and OsTIR1 (F74G)-V5 Kpn1 Reverse (GATGGTACCTCACGTAGAATCGAGACCGA) primers.

OsTIR1 (F74G)-V5 PCR product was purified using the QIAquick PCR Purification Kit (QIAGEN) following the standard protocol. To generate OsTIR1 (F74G)-V5 in pENTR1A, the OsTIR1 (F74G)-V5 PCR product was subcloned into the pENTR1A no ccDB (w48-1) entry vector using Sal1 and KpnI restriction sites. The OsTIR1 (F74G)-V5 in pENTR1A was then recombined with the pLenti CMV-Neo-DEST (705-1) vector under the CMV promoter using Gateway LR Clonase II Enzyme Mix according to the manufacturer’s instructions. The OsTIR1 (F74G)-V5 lentiviral particles were prepared as described previously. This lentivirus was used to transduce hTERT-RPE1 VPS54-KO cells expressing mVPS54-mAID. The transduced cells were tested for mVPS54-mAID/OsTIR1 (F74G)-V5 co-expression by WB and IF. Single-cell clones were then isolated by serial dilution, expanded, and characterized. To induce rapid VPS54 depletion in resulting cells, the auxin analog 5-phenyl-indole-3-acetic acid (5-Ph-IAA) (10 µM) was added at various time points. For convenience, auxin analog 5-phenyl-indole-3-acetic acid (5-Ph-IAA) will be named as AA hereafter.

HeLa VPS54-mAID OsTIR1 (F74G)-V5 cells were generated using a slightly different procedure. Briefly, HeLa VPS54-KO cells were co-transfected with AAVS1-Tet-OsTIR1 (F74G)-V5 and AAVS1 T2 CRISPR in pX330. After 48 hours, selection was performed with 2 µg/ml puromycin. Single-cell sorting was conducted to isolate OsTIR1 (F74G)-V5 positive clones. Once these clones were established, they were transduced with mVPS54-mAID lentiviruses. Single-cell clones expressing mVPS54-mAID OsTIR1 (F74G)-V5 were isolated by serial dilution. Expression of OsTIR1 (F74G)-V5 was induced by doxycycline (2 µg/ml) for 24 hours before the experiment. To induce rapid VPS54 depletion in HeLa cells, the auxin analog AA (10 µM) was added at various time points.

### Construction of RPE1 cell lines stably expressing MPR-mNeonGreen and mScarlet-GS15

MPR-mNeonGreen-P2A-mScarlet3-GS15-pUC57 construct synthesized by Genescript initially subcloned into pENTR1A using BamH1 and XhoI sites. This construct was then recombined into the pLenti-COG4_pr_-Neo-DEST plasmid, using Gateway LR Clonase II Enzyme Mix (Thermo Fisher). The recombined plasmid was transformed into Stbl3 competent cells as per the manufacturer’s instructions, and DNA was extracted using the QIAprep Spin Miniprep Kit. Correct MPR-mNG-P2A-mScarlet-GS15 pLenti clones were verified by restriction analysis. The expression of MPR-mNG and mScarlet-GS15 was validated by WB and IF analysis of transfected HEK293T cells. The MPR-mNG-P2A-mScarlet-GS15 lentivirus was prepared as described previously and RPE1 mVPS54-mAID expressing cells were transduced and sorted for single cell clones.

### Preparation of cell lysates and Western blot analysis

For preparation of cell lysates, cells grown on tissue culture dishes were washed twice with PBS and lysed in 2% SDS that was heated for 5 min at 70°C. Total protein concentration in the cell lysates was measured using the BCA protein assay (Pierce). The protein samples were prepared in 6X SDS sample buffer containing beta-mercaptoethanol and denatured by incubation at 70°C for 10 minutes. 10-30 µg of protein samples were loaded onto Bio-Rad (4-15%) gradient gels or Genescript (8-16%) gradient gels. Gels were transferred onto nitrocellulose membranes using the Thermo Scientific Pierce G2 Fast Blotter. Membranes were rinsed in PBS, blocked in Odyssey blocking buffer (LI-COR) for 20 min, and incubated with primary antibodies overnight at 4°C. Membranes were washed with PBS and incubated with secondary fluorescently tagged antibodies diluted in Odyssey blocking buffer for 60 min. Blots were then washed and imaged using the Odyssey Imaging System. Images were processed using the LI-COR Image Studio software. Primary and secondary antibodies used in this work are listed in Table 2.

**Table 2.**
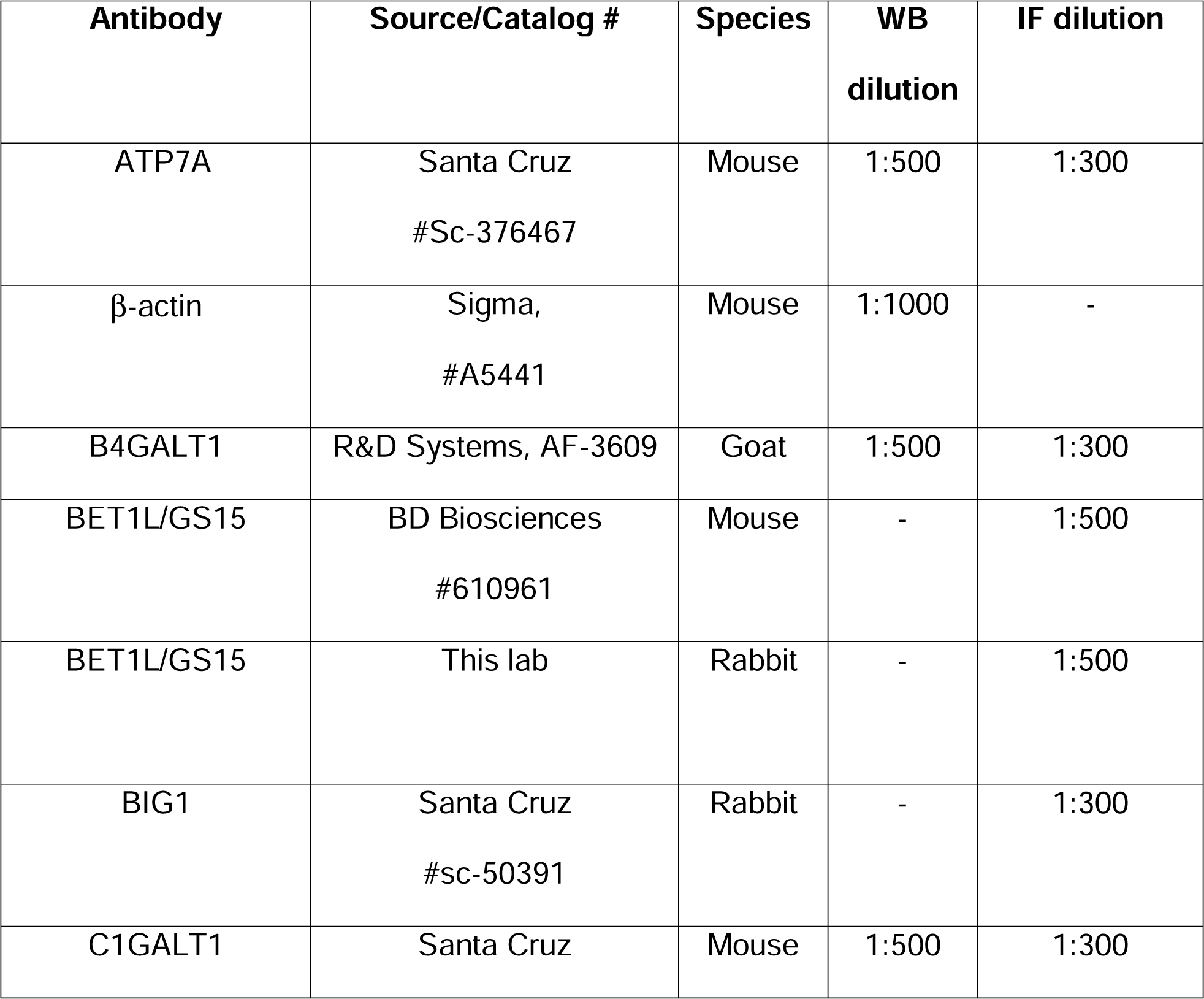
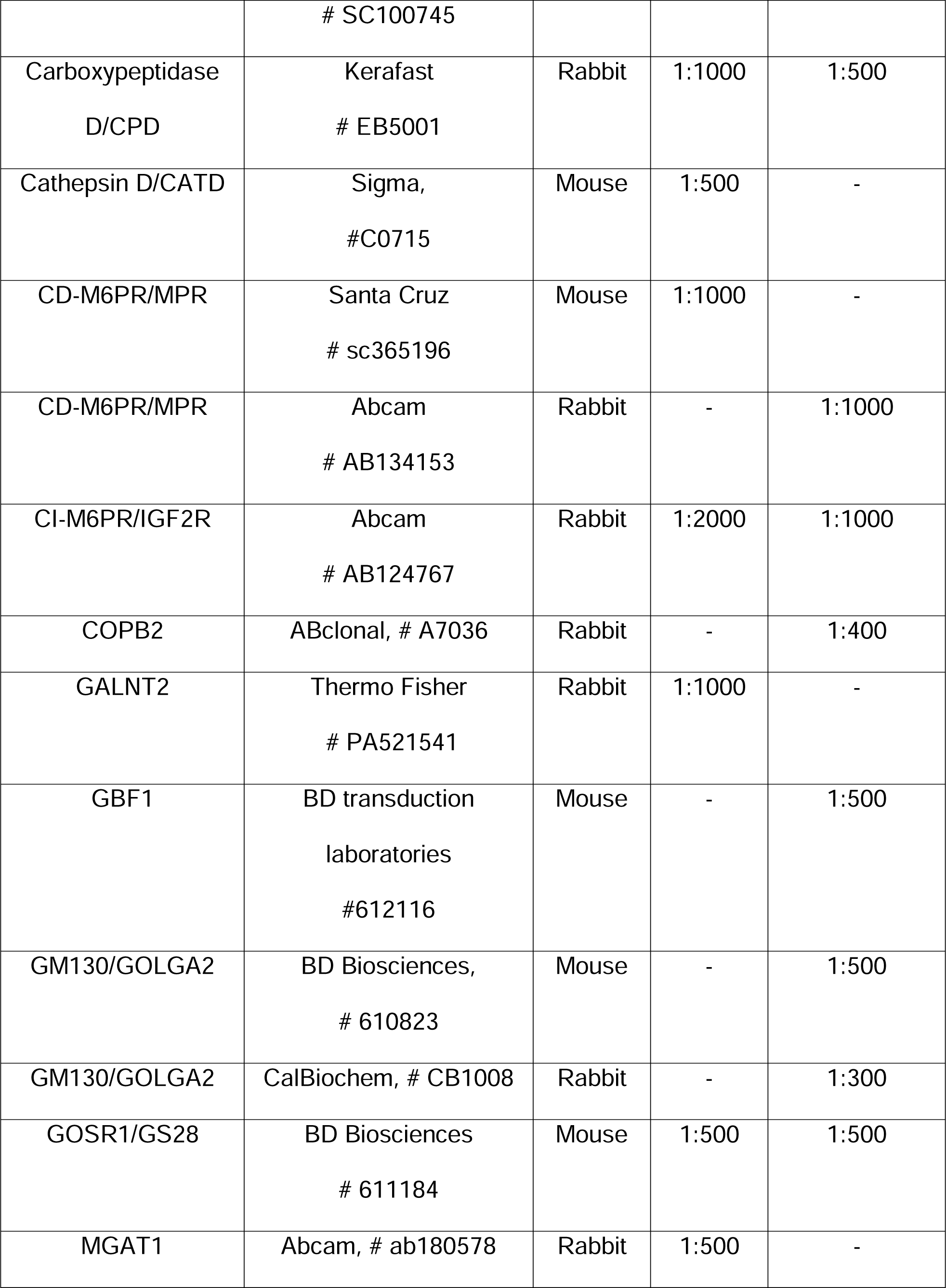
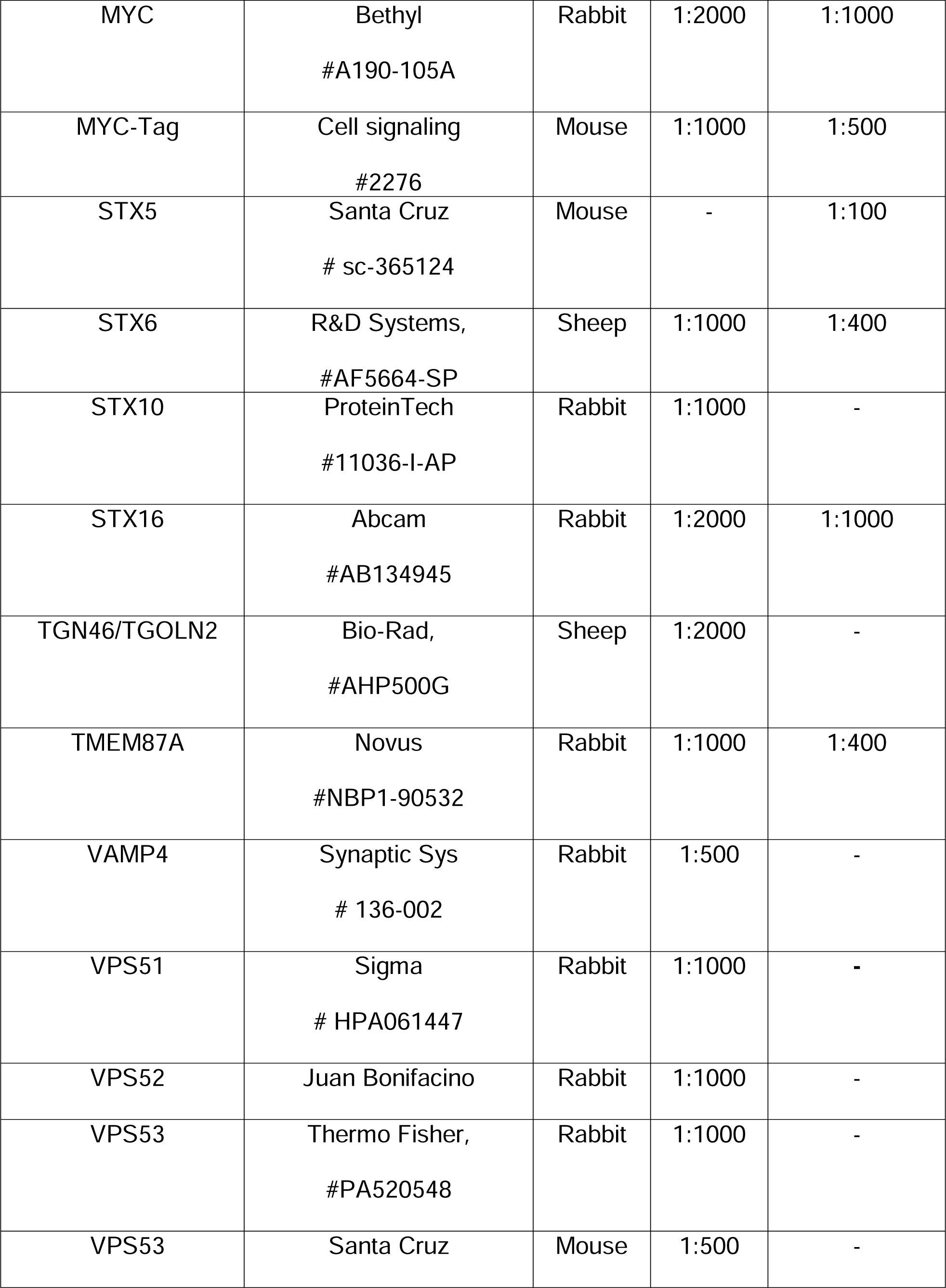
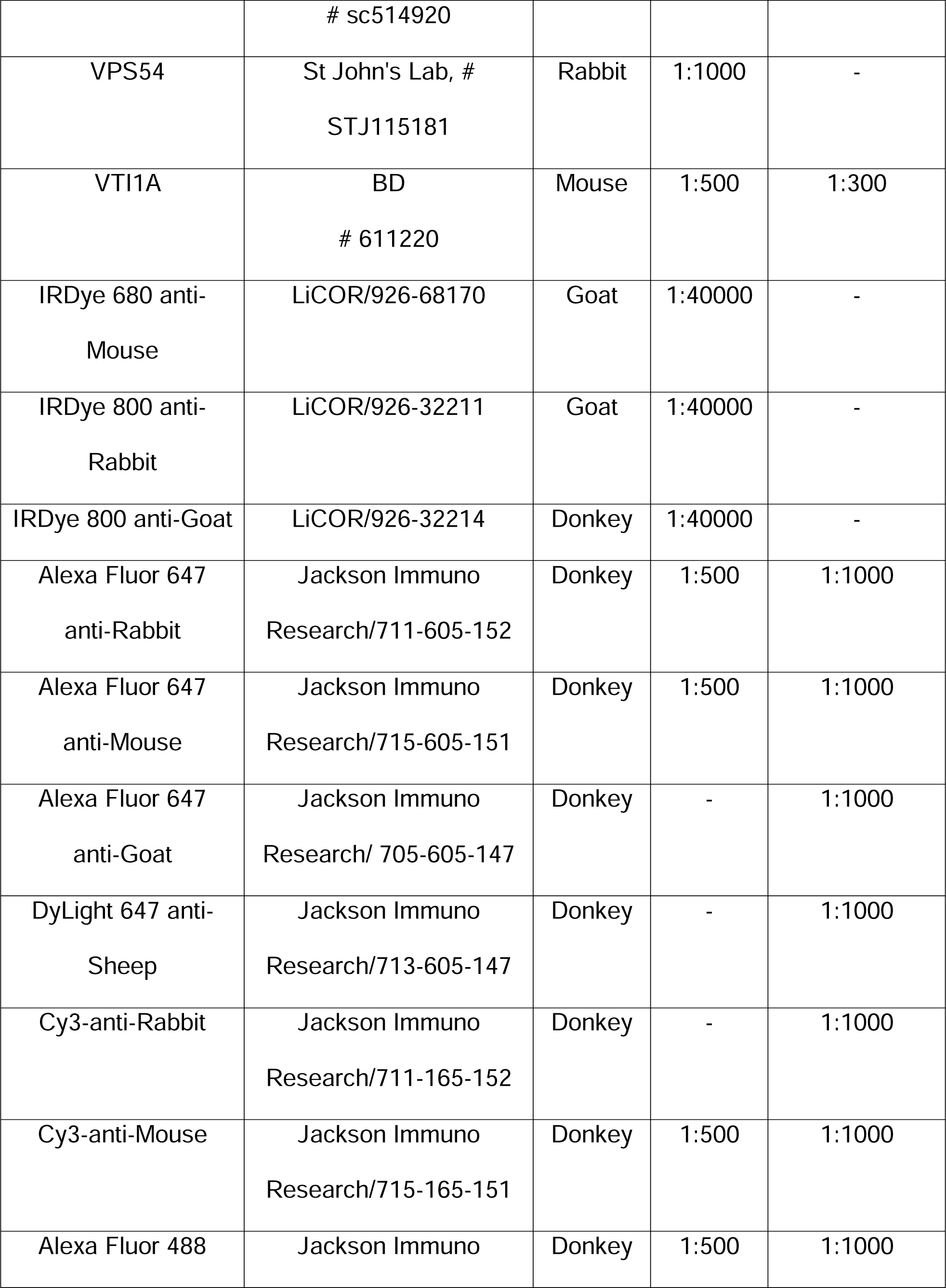
List of primary and secondary antibodies.

### Lectin blotting and staining

To perform blots with fluorescent lectins, 10 µg of cell lysates were loaded onto Bio-Rad (4-15%) gradient gels and run at 160V. Next, proteins were transferred to nitrocellulose membrane using the Thermo Scientific Pierce G2 Fast Blotter. The nitrocellulose membrane was blocked with 3% bovine serum albumin (BSA) for 30 minutes. The lectins Helix Pomatia Agglutinin (HPA) or Galanthus Nivalis Lectin (GNL) conjugated to Alexa 647 fluorophore were diluted 1:1000 in 3% BSA from their stock concentration of 1 µg/µl and 5 µg/µl, respectively. Blots were incubated with lectin solutions for 30 min and then washed in PBS four times for four minutes each and imaged using the Odyssey Imaging System.

### Immunofluorescence microscopy

Cells were plated on glass coverslips to 80-90% confluency and fixed with 4% paraformaldehyde (PFA) (freshly made from 16% stock solution) in phosphate-buffered saline (PBS) for 15 minutes at room temperature. Cells were then permeabilized with 0.1% Triton X-100 for one minute followed by treatment with 50 mM ammonium chloride for 5 minutes and washed with PBS. After washing and blocking twice with 1% BSA, 0.1% saponin in PBS for 10 minutes, cells were incubated with primary antibody (diluted in 1% cold fish gelatin, 0.1% saponin in PBS) for 40 minutes, washed, and incubated with fluorescently conjugated secondary antibodies for 30 minutes. Cells were washed four times with PBS, then coverslips were dipped in PBS and water 10 times each and mounted on glass microscope slides using Prolong® Gold antifade reagent (Life Technologies). Cells were imaged with a 63× oil 1.4 numerical aperture (NA) objective of a LSM880 Zeiss Laser inverted microscope and Airyscan super resolution microscope using ZEN software. Quantitative analysis was performed using single-slice confocal images. All the microscopic images shown are Z-stacked Maximum Intensity Projection images.

### Live cell microscopy

Cells were plated on 35 mm glass bottom dishes with No. 1.5 coverglass (MatTek Corporation). Transfection was performed using Lipofectamine 3000. After 16–18 hours, just before imaging, the media was replaced with warm FluoroBrite™ DMEM Media (Gibco, Cat # A1896701) supplemented with 10% FBS. Imaging was conducted on an LSM880 Zeiss inverted microscope equipped with confocal optics, using a 63× oil objective with a 1.4 numerical aperture (NA) and Airyscan. During imaging, the environment was maintained at 37°C, 5% CO2, and 90% humidity.

### Cell fractionation

Cells grown to 90% confluency in 15 cm dishes were washed with PBS and collected by trypsinization, followed by centrifugation at 400×g for 5 minutes. The cell pellet was resuspended in 1.5 ml of cell collection solution (0.25 M sucrose in PBS) and centrifuged again at 400×g for 5 minutes. The pellet was then resuspended in 1.5 ml of hypotonic lysis solution (20 mM HEPES, pH 7.2, with a protein inhibitor cocktail and 1 mM PMSF) and passed through a 25 G needle 20 times to lyse the cells. Cell lysis efficiency was assessed under a phase-contrast microscope. Subsequently, KCl (to a final concentration of 150 mM) and EDTA (to a final concentration of 2 mM) were added. Unlysed cells and nuclei were removed by centrifugation at 1000×g. The post nuclear supernatant (PNS) was transferred to a 1.5 ml Beckman tube, and the Golgi-enriched fraction (P30) was pelleted by centrifugation at 30,000×g for 10 minutes. The supernatant (S30) was then transferred to a new Beckman tube, and the vesicle-enriched fraction was isolated by centrifugation at 100,000×g for 1 hour at 4°C using a TLA-55 rotor.

### Vesicle Immunoprecipitation (GS15 IP)

Cells grown to 90% confluency in 15 cm dishes were washed with PBS and collected by trypsinization, followed by centrifugation at 400×g for 5 minutes. The cell pellet was resuspended in 1.5 ml of cell collection solution (0.25 M sucrose in PBS) and centrifuged again at 400×g for 5 minutes. The pellet was then resuspended in 1.5 ml of hypotonic lysis solution (20 mM HEPES, pH 7.2, with a protein inhibitor cocktail and 1 mM PMSF) and passed through a 25 G needle 20 times to lyse the cells. Cell lysis efficiency was assessed under a phase-contrast microscope. Subsequently, KCl (to a final concentration of 150 mM) and EDTA (to a final concentration of 2 mM) were added. Unlysed cells and nuclei were removed by centrifugation at 1000×g. The postnuclear supernatant (PNS) was transferred to a 1.5 ml Beckman tube, and the Golgi-enriched fraction (P30) was pelleted by centrifugation at 30,000×g for 10 minutes. The supernatant (S30) was transferred to a new tube containing 10 µl of GS15 antibody and incubated at room temperature on a rotating platform for 2 hours. Subsequently, 30 µl of Dyna Protein G magnetic beads (ThermoFisher Scientific #10004D) were added to the tube with the S30 and GS15 antibody mixture. This mixture was rotated at room temperature for an additional 1 hour. The protein bound to the beads were eluted by adding 2x sample buffer with 10% β-mercaptoethanol and heated at 95°C in a heat block for 5 min.

### Secretion assay

hTERT-RPE1-VPS54-mAID expressing cells were plated in three 6-cm dishes and grown to 90-100% confluency. Cells were then rinsed 3 times with PBS and placed in 2 ml serum-free, chemically defined medium (BioWhittaker Pro293a-CDM, Lonza) with 1× GlutaMAX (100× stock, Gibco) added per well for 48 hours. 42 hours post-incubation of cells in serum-free, chemically defined medium, one of the wells was treated with 10 µM of AA and the other well was used as control. After completion of 48 hours incubation, the supernatant was collected and spun down at 3,000xg to remove floating cells. The supernatant was concentrated using a 10k concentrator (Amicon® Ultra 10k, Millipore); final concentration was 10× that of cell lysates.

### High-pressure freezing, freeze substitution, and Electron Microscopy

Sapphire disks were initially coated with a 10 nm carbon layer, followed by a collagen (Corning) coating according to the manufacturer’s protocol. The coated disks were sterilized under UV light and transferred into new sterile 3 cm dishes for plating the cells. After the cells reached 80%–100% confluence, they were incubated in fresh media for 2–3 hours at 37°C to equilibrate, then treated with Auxin for 0 hour and 3 hours respectively. High-pressure freezing (HPF) was carried out at designated time points in a cryo-protectant solution (PBS with 2% Type IX ultra-low melt agarose (Sigma-Aldrich), 100 mM D-mannitol, and 2% FBS). This procedure used a Leica EM PACT2 high-pressure freezing unit (Leica Microsystems) equipped with a rapid transfer system, maintaining a high-pressure of 2100 bar. All solutions, bayonets, and sample holders were pre-warmed to 37°C, and every step of the process was performed on a 37°C heating platform to ensure consistent temperature control.

### Freeze substitution dehydration

Samples were transferred under liquid nitrogen into cryovials containing anhydrous acetone with 2% osmium tetroxide (OsO4), 0.1% glutaraldehyde, and 1% double-distilled (dd) H_2_O. The cryovials were then placed into a freeze-substitution chamber set at −90°C and subjected to the following schedule: maintained at −90°C for 22 hours, warmed at 3°C per hour to −60°C, held at −60°C for 8 hours, then warmed at 3°C per hour to −30°C, and kept at −30°C for 8 hours before warming to 0°C. Afterward, the samples were placed on ice and transferred to a cold room set at 4°C. Following three washes with acetone, the samples were stained with a solution of 1% tannic acid and 1% ddH_2_O in acetone on ice for 1 hour, followed by another three acetone washes.

Next, the samples were stained with a 1% OsO4 and 1% ddH_2_O solution in acetone on ice for 1 hour. Afterward, they were washed three times for 10 minutes each in acetone and dehydrated through a graded ethanol series (25%, 50%, 75%, and 100%) using automatic resin infiltration. protocol for PELCO Bio-Wave Pro laboratory microwave system. Samples were embedded in Araldite 502/Embed 812 resins with a DMP-30 activator and baked at 60°C for 48 h.

### Thin section TEM

Thin sections, 50 nm in thickness, were cut using a Leica UltraCut-UCT microtome and subsequently post-stained with aqueous uranyl acetate and Reynold’s lead citrate (EMS).

### Electron microscopy and image handling

Images were taken using an FEI Tecnai TF20 intermediate-voltage electron microscope operated at 80 keV (FEI Co.). The images were acquired with an FEI Eagle 4 k digital camera controlled with FEI software.

### Colocalization analysis

Pearson’s correlation coefficient was calculated using “Colocalization” module of Zen Blue software. The colocalization between different proteins was recorded and the graph was made using GraphPad Prism 9.3.0. At least 30 cells were used for quantification of Golgi area per group and Pearson’s correlation coefficient was measured.

### Statistical analysis

All results are representative of at least 3 independent experiments. Western blot images are representative from 3 repeats. Western blots were quantified by densitometry using the LI-COR Image Studio software. Error bars for all graphs represent standard deviation. Statistical analysis was done using one-way ANOVA, two-way ANOVA or paired t test using GraphPad Prism software.

## Results

### Development of the rapid GARP inactivation system

Previous investigation of hTERT-RPE1 GARP-KO cells [36] [46] [35] revealed that VPS54-KO specifically inactivates GARP complex, resulting in dramatic changes in Golgi structure and function. To uncover primary defects associated with GARP dysfunction, an auxin-inducible degron version 2 (AID2) system [37] was utilized. VPS54, the unique subunit of the GARP complex, was tagged with plant degron mAID and stably expressed under the control of the COG4 promoter region [47] in the RPE1 VPS54-KO cells. The constructed cellular system also expressed auxin receptor OsTIR1 (F74G) mutant that, in the presence of auxin homolog 5-phenyl-indole-3-acetic acid (AA) should form a complex with mAID, directing the hybrid protein for poly-ubiquitination and proteasomal degradation. First, we tested the functionality of VPS54-mAID protein by western blot (WB) (Figure 1 A, B) and immunofluorescence microscopy (IF) analysis (Figure 1C). We found that a decrease in total protein abundance of TGN46/TGOLN2, B4GALT1, and GS15/BET1L observed in VPS54-KO cells was restored upon the expression of VPS54-mAID (Figure 1C). Furthermore, a decrease in colocalization of TGN46 (Figure 1D) and GS15 (Figure 1E) with the *trans*-Golgi marker P230/GOLGA4 in VPS54-KO cells was rescued in VPS54-KO cells expressing VPS54-mAID. A similar functionality test of VPS54-mAID in HeLa VPS54-KO cells revealed that proper Golgi localization of TGN46, GBF1, and COPB2 was restored upon expression of VPS54-mAID (Figure S1A-C). Hence, the VPS54-mAID construct is functional.

**Figure 1:**
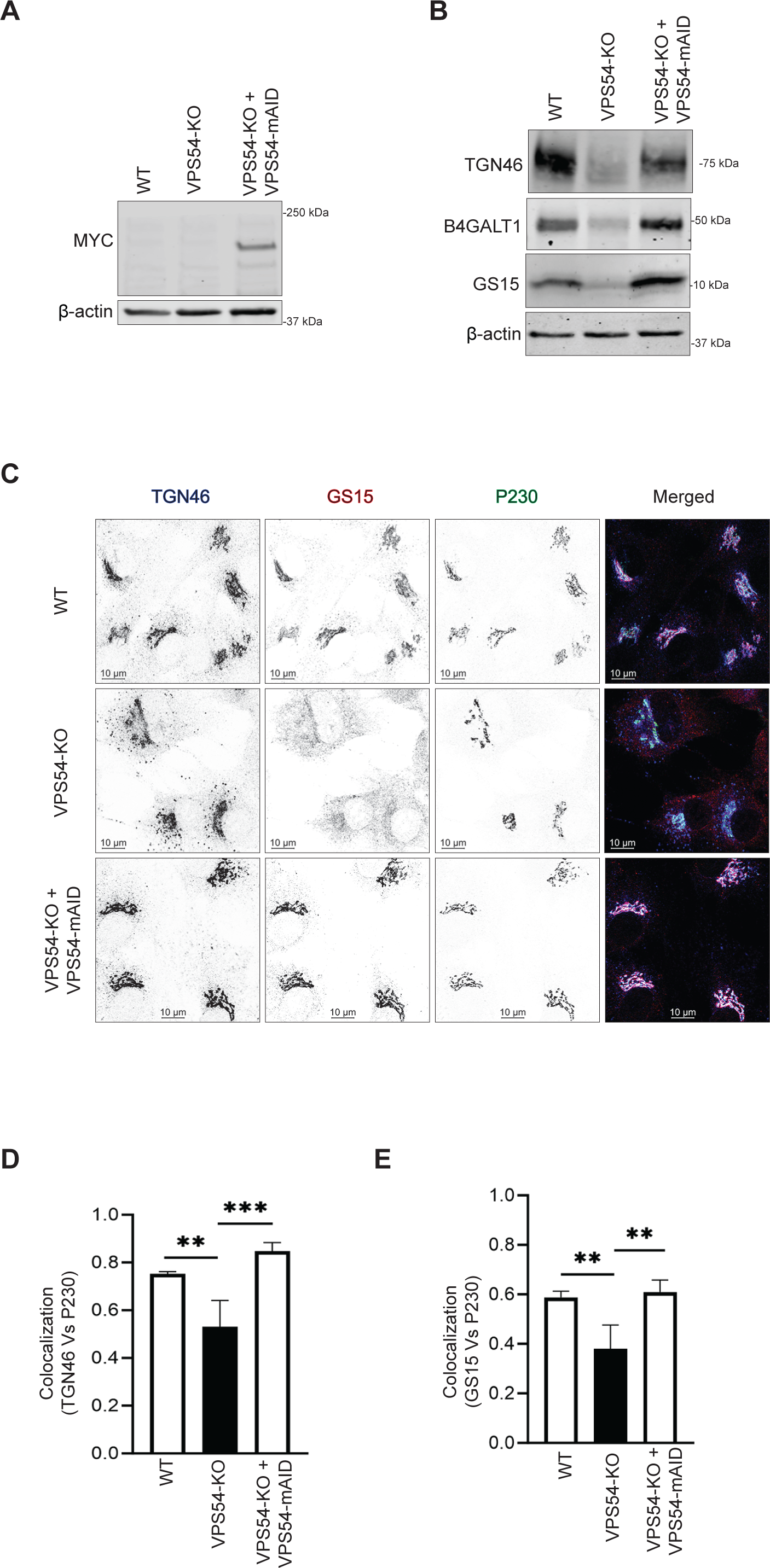
Expression of VPS54-mAID rescues VPS54-KO defects. (A) Western blot (WB) analysis of RPE1 cell lysates from wild-type (WT), VPS54 knock out (VPS54-KO), and VPS54-KO cells rescued with VPS54-mAID. Blots were probed with anti-myc (to detect VPS54-myc-mAID) and anti-β-actin antibodies. (B) WB analysis of RPE1 cell lysates from WT, VPS54-KO, and VPS54-mAID, probed with anti-TGN46, anti-B4GALT1, and anti-GS15 antibodies. β-actin was used as a loading control. (C) Confocal microscopy images of WT, VPS54-KO, and VPS54-mAID RPE1 probed for TGN46, GS15, and P230. (D) Quantification of IF images in (C). Pearson’s correlation coefficient was used to assess the colocalization of TGN46 and P230. (E) Pearson’s correlation coefficient was used to assess the colocalization of GS15 and P230. At least 50 cells were analyzed per sample for the quantification. Statistical significance was determined using one-way ANOVA. ** p≤ 0.01, *** p ≤ 0.001.

### Acute depletion of VPS54 does not affect the protein abundance of its protein partners

Once we confirmed that the cells expressing VPS54-mAID could rescue the VPS54-KO defects, we next aimed to induce the rapid depletion of VPS54 by treating the cells with 5-Ph-IAA (AA, Figure 2A). We tested the efficiency of VPS54 depletion by treating the cells with AA for 0, 0.5, 1, 2, and 3 h, respectively. WB and IF analysis demonstrated that approximately 70 % of the VPS54 was depleted in 30 minutes, and in 3 hours, almost all VPS54 was degraded (Figure 2B-C). Prolonged (24-48 h) treatment with AA resulted in a continuous depletion of VPS54-mAID (data not shown). A similar rapid depletion of VPS54 was observed in HeLa VPS54-KO cells expressing VPS54-mAID (Figure S2A). IF analysis confirmed a complete depletion of VPS54-mAID in the Golgi of AA-treated cells (Figure S2B). We next examined if the depletion of VPS54 can lead to the degradation of other GARP subunits. In agreement with the data obtained with VPS54-KO cells (unpublished data), the total protein abundance of VPS51, VPS52, and VPS53 remains mostly unchanged in cells acutely depleted for VPS54 (Figure 2D-F). Their unchanged protein abundance indicates that rapid VPS54 depletion has not resulted in destabilization and degradation of the EARP complex, as VPS51, VPS52, and VPS53 are the shared subunits of GARP and EARP complexes.

**Figure 2:**
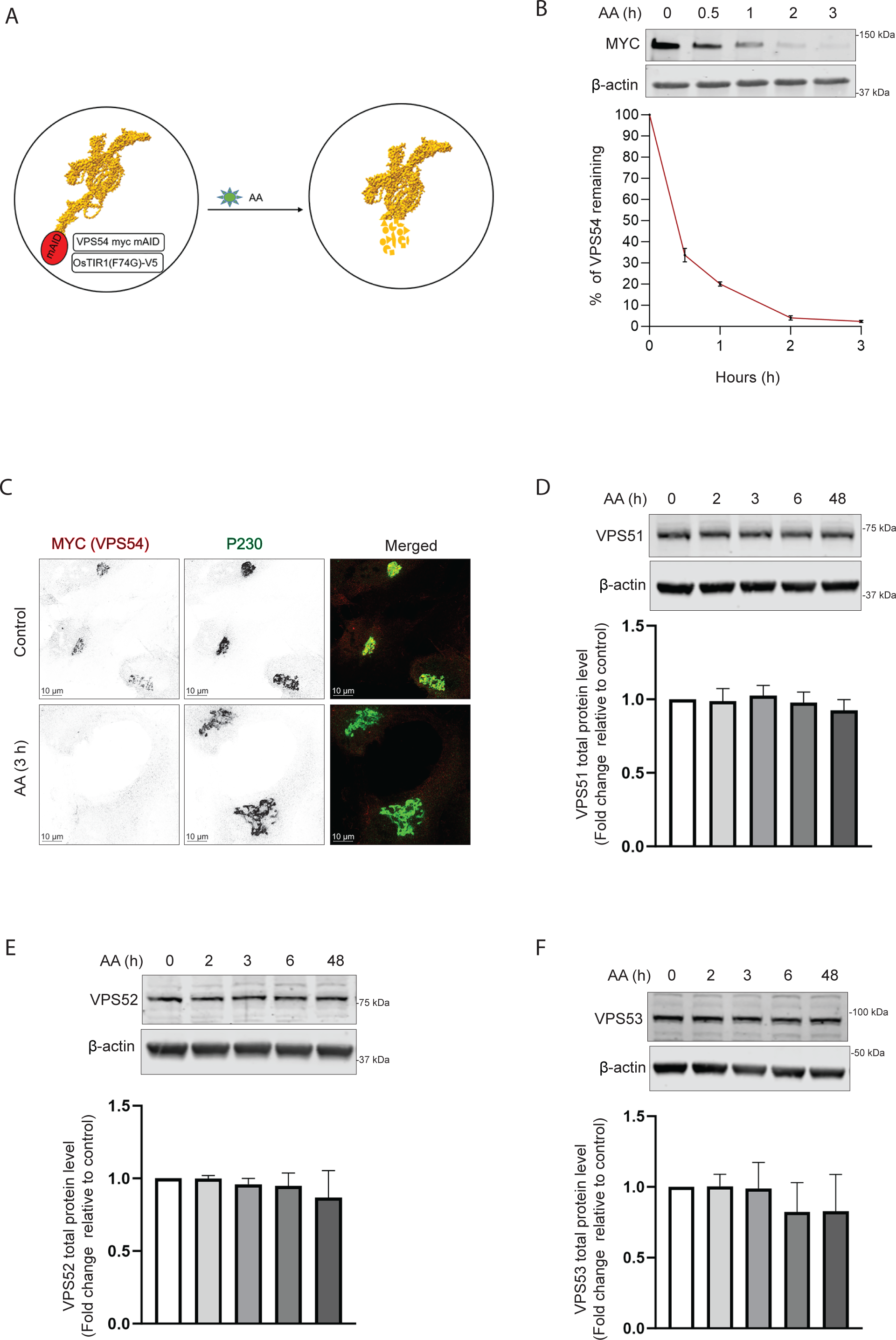
VPS54-mAID rapid depletion does not affect the stability of other GARP subunits. (A) Diagram illustrating the cellular setup for the rapid depletion of VPS54 subunit of GARP complex using 5-Ph-IAA (Auxin Analogue or AA). (B) RPE1 cells expressing VPS54-mAID were treated with AA for 0, 0.5, 1, 2, and 3 h respectively to deplete VPS54. (Top panel) WB with anti-myc antibody. (Bottom Panel) Quantification of the blots from three independent experiments. (C) RPE1 cells expressing VPS54-mAID were treated with AA for 3 h and co-stained for myc (red) and P230 (green). (D) WB of RPE1 cells expressing VPS54-mAID, treated with AA for 0, 2, 3, 6, and 48 h was performed using anti-VPS51 antibody. (E) WB of RPE1 cells expressing VPS54-mAID, treated with AA for 0, 2, 3, 6, and 48 h was performed using anti-VPS52 antibody. (F) WB of RPE1 cells expressing VPS54-mAID, treated with AA for 0, 2, 3, 6, and 48 h was performed using anti-VPS53 antibody. The bottom panels in (D), (E), and (F) show quantification of the blots from three independent experiments.

### Acute depletion of VPS54 alters the protein abundance and localization of a subset of TGN proteins

The GARP complex is believed to tether the endosome-derived vesicles at the TGN. Several TGN resident proteins, including TGN46/TGOLN2, ATP7A, TMEM87A, CPD, and mannose-6-phosphate receptors, are known to cycle between the endosomes and Golgi [48] [32].

TGN46 is a single-pass type I transmembrane protein believed to function as a receptor for secretory cargoe [49]. TGN46 is localized to the TGN in a steady state, it cycles between the TGN, endosomes, and the PM [50] [51] [52] [53] [54]. Since the TGN46 was significantly depleted in GARP-KO cells [36], we reasoned that TGN46 instability could be a primary defect of GARP dysfunction. Indeed, we found that TGN46 was significantly depleted within 3 hours of the induction of VPS54 degradation (Figure 3A). Additionally, TGN46 was significantly mislocalized from the Golgi to peripheral punctate structures in VPS54-depleted cells. TGN46 mislocalization was specific since the localization of non-cycling peripheral membrane proteins, such as the golgins GM130/GOLGA2 and P230/GOLGA4, was unaffected by GARP dysfunction (Figure 3B, E). Indeed, VPS54 depletion resulted in a significant decrease in colocalization of TGN46 with P230 (Figure 3C). As discussed later, it’s possible that following the rapid degradation of VPS54, the TGN46 is rerouted to endolysosomes for lysosomal degradation. In support of this model, treating VPS54-depleted cells with lysosomal protease inhibitor (PI) resulted in partial restoration of TGN46 expression (Figure S3A). Furthermore, co-transfection of VPS54-depleted cells with rat homolog of TGN46, TGN38-GFP, and endolysosomal marker Lamp2-mCherry resulted in partial colocalization of TGN38 with lysosomes (Figure S3B).

**Figure 3:**
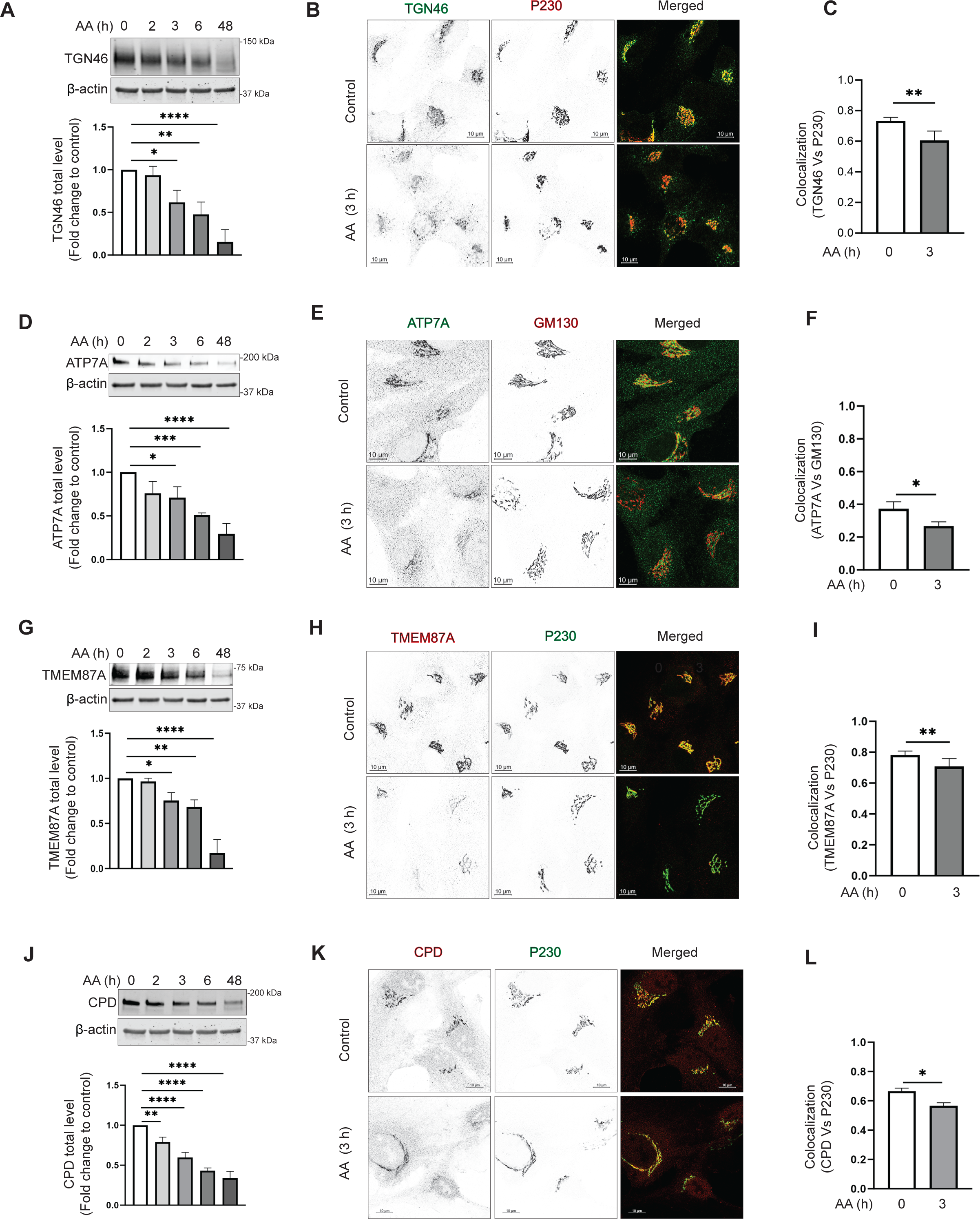
Acute depletion of VPS54 alters the abundance and localization of TGN proteins. (A), (D), (G), and (J) RPE1 cells expressing VPS54-mAID were treated with AA for 0, 2, 3, 6, and 48 h respectively to deplete VPS54. (Top panels) WB analysis of cell lysates was performed and probed with (A) anti-TGN46, (D) anti-ATP7A, (G) anti-TMEM87A, and (J) anti-CPD antibody. (Bottom panels) Quantification of the blots from three independent experiments. (B), (E), (H), (K) RPE1 cells expressing VPS54-mAID were treated with AA for 3 h and co-stained for (B) TGN46 and P230, (E) ATP7A and GM130, (H) TMEM87A and P230, (K) CPD and P230. (C), (F), (I), and (L) Colocalization analysis was performed by calculating the Pearson’s correlation coefficient for (C) TGN46 and P230, (F) ATP7A and GM130, (I) TMEM87A and P230, and (L) CPD and P230. At least 50 cells were imaged per sample for the quantification. Statistical significance was assessed using one-way ANOVA. ** p≤ 0.01, * p ≤ 0.05.

Menkes proteins (also known as ATP7A/B) are integral to the mammalian copper transport system, cycling continuously between the Golgi complex and the plasma membrane [55] [56]. VPS54-KO resulted in mislocalization of ATP7A that was reversed by expression of VPS54-mAID, indicating that ATP7A cycling is GARP-dependent (Figure S3C). Indeed, the total protein abundance of ATP7A was significantly decreased within 3 hours of the induction of VPS54 degradation (Figure 3D). Consistent with ATP7A mislocalization in GARP-KO cells, acute depletion of VPS54 also altered Golgi localization of ATP7A (Figure 3E, F). The internal environment of the Golgi is slightly acidic at pH 6.0-6.7 and is maintained by ion channels such as Golgi-pH-regulating cation channel GolpHCat/TMEM87A [57]. We have discovered a significant decrease in the total protein level of TMEM87A in VPS54-depleted cells, indicating that TMEM87A recycling depends on GARP function (Figure 3G). In agreement to TMEM87A sensitivity in GARP-KO cells, the Golgi localization of TMEM87A and protein stability was significantly decreased in cells acutely depleted for VPS54 (Figure 3H-I).

Carboxypeptidase D/CPD, a transmembrane TGN enzyme is known to recycle through endosomes and the plasma membrane [58]. IF analysis of CPD localization in wild-type and VPS54-KO RPE1 cells confirmed its Golgi localization and revealed a decrease in Golgi staining in GARP-KO cells (Figure S3D). CPD stability and localization were significantly affected in cells acutely depleted for VPS54 (Figure 3J-L). Hence, the stability and localization of four TGN transmembrane proteins was specifically altered upon rapid GARP inactivation.

### Rapid VPS54 depletion causes Cathepsin D sorting defects and enhances fibronectin secretion, without significant alterations in stability or localization of mannose-6-phosphate receptors

MPRs (mannose-6-phosphate receptors) are crucial for transporting lysosomal enzymes, like Cathepsin D, from the Golgi to the endosomes and then to the lysosomes [59]. There are two types of MPRs: cation-dependent MPR (CD-MPR/MPR) and cation-independent MPR (CI-MPR/IGF2R) [60] [61] [48]. After delivering their cargo, MPRs are recycled back to the Golgi for subsequent rounds of enzyme transport, and GARP is expected to be a part of the recycling machinery for MPRs [11]. Our previous work on GARP-KO cells showed an increase in the secretion of pro-Cathepsin D [36]. In agreement with the data in VPS54-KO cells, we observed a significant increase in the secretion of pro-Cathepsin D from cells acutely depleted for VPS54 (Figure 4A). At the same time, no changes in intracellular mature Cathepsin D or its precursor were observed (Figure 4B). Interestingly, pro-Cathepsin D secretion was accompanied by the increased secretion of Fibronectin/FN1 (Figure 4C), indicating a dysfunction of TGN protein sorting machinery in cells acutely depleted for VPS54. We further investigated if GARP dysfunction stimulates the fibronectin release or if this is a result of protein overproduction and found that the intracellular fibronectin level remains unchanged (Figure 4D). These results collectively indicate that VPS54 acute depletion leads to TGN sorting defects.

**Figure 4:**
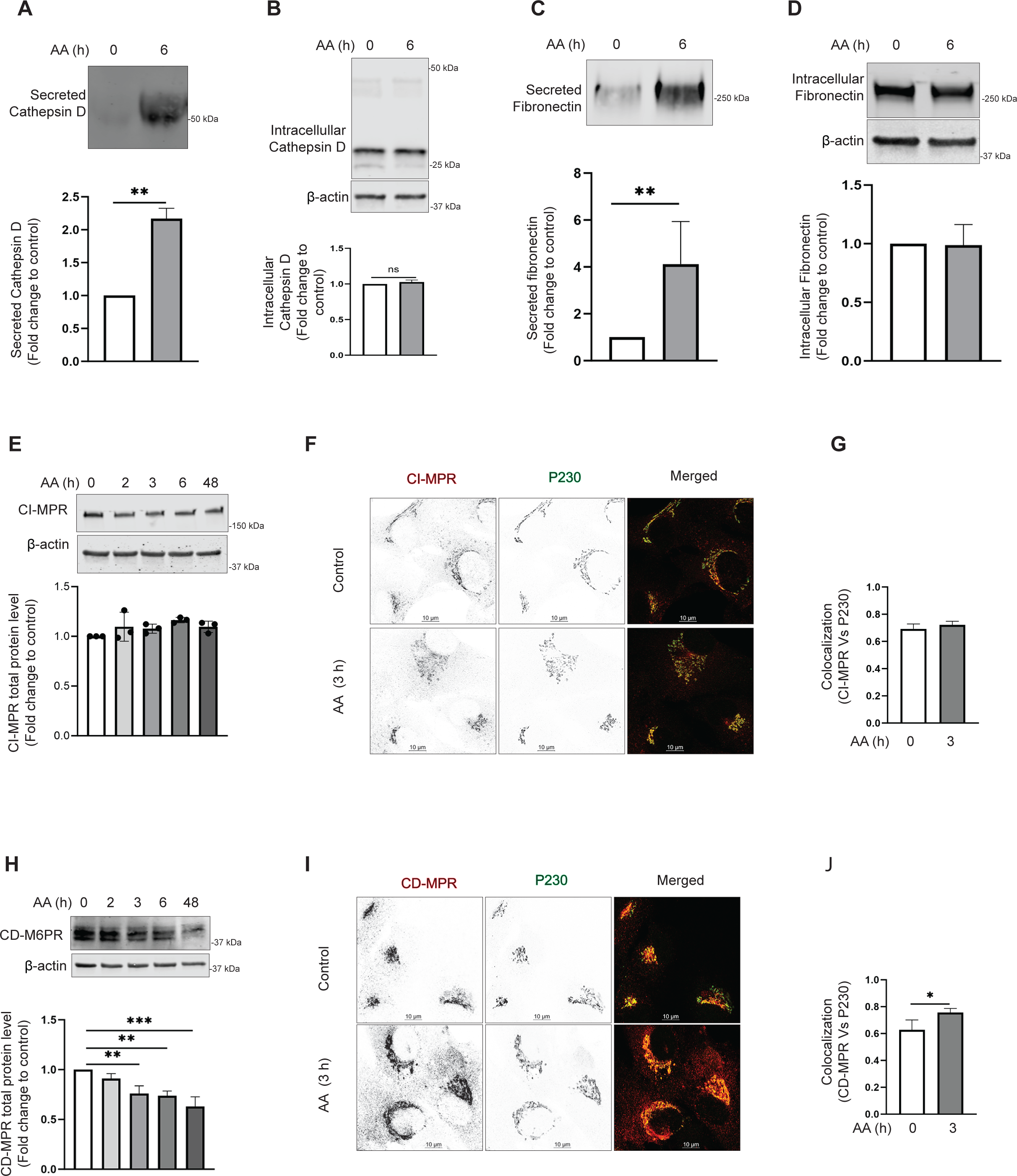
Acute VPS54 depletion causes secretory defects and relocalization of CD-MPR to vesicle. (A) (Top panel) WB analysis of secreted Cathepsin D from RPE1 VPS54-mAID cells treated with AA for 0 h and 6 h respectively. (Bottom panel) Quantification of secreted Cathepsin D from three independent experiments. (B) WB analysis of the whole cell lysates from (A), probed with anti-Cathepsin D antibody. (Bottom panel) Quantification of intracellular Cathepsin D from three independent experiments. (C) (Top panel) WB analysis of the secreted Fibronectin from RPE1 VPS54-mAID cells treated with AA for 0 h and 6 h respectively. (Bottom panel) Quantification of secreted Fibronectin from three independent experiments. (D) WB analysis of the whole cell lysates from (C), probed with anti-Fibronectin antibody. (Bottom panel) Quantification of intracellular Fibronectin from three independent experiments. Statistical significance was assessed using paired t-test. ** p≤ 0.01. (E) (Top panel) WB analysis of RPE1 cells expressing VPS54-mAID, treated with AA for 0, 2, 3, 6, and 48 h respectively and probed with anti-CI-MPR antibody. (Bottom panel) Quantification of the blots from three independent experiments. (F) RPE1 VPS54-mAID expressing cells were treated with AA for 3 h and co-stained for CI-MPR and P230. (G) Colocalization of CI-MPR to P230 was calculated between control and 3 h AA treatment groups using Pearson’s correlation coefficient and graph was prepared in GraphPad prism. (H) (Top panel) WB analysis of RPE1 cells expressing VPS54-mAID treated with AA for 0, 2, 3, 6, and 48 h respectively and probed with anti-CD-MPR antibody. (Bottom panel) Quantification of the blots from three independent experiments. (I) RPE1 VPS54-mAID expressing cells were treated with AA for 3 h and cells were co-stained for CD-MPR and P230. (J) Colocalization of CD-MPR to P230 was calculated between the control and 3 h AA treatment groups using Pearson’s correlation coefficient and graph was prepared in GraphPad prism. Statistical significance was calculated using paired t-test. * p≤ 0.05

Previous investigation of MPRs localization in HeLa cells suggested that siRNA depletion of VPS52 resulted in “accumulation of recycling MPRs in a population of light, small vesicles downstream of endosomes” [11]. To test if this is the case in cells rapidly depleted for VPS54, the stability and localization of CD-MPR and CI-MPR was tested. The total protein level of CI-MPR and its Golgi localization were not significantly changed between control and VPS54-depleted cells (Figure 4E-G), indicating that the trafficking pathway and/or machinery of CI-MPR are different from other TGN transmembrane proteins. WB analysis showed that the protein level of CD-MPR significantly decreased following acute VPS54 depletion (Figure 4H), coinciding with the appearance of a vesicle-like haze surrounding the Golgi (Figure 4I). Interestingly, the Pearson coefficient of colocalization between CD-MPR and the TGN marker golgin P230 increased in VPS54-depleted cells (Figure 4J), indicating that CD-MPR responds to GARP depletion in a manner distinct from other TGN resident proteins. This suggests that CD-MPR may follow a unique trafficking or retention pathway under GARP-deficient conditions. The data suggests that the missorting of cathepsin D in GARP-depleted cells is possibly unrelated to the mistargeting of MPRs.

### Acute GARP dysfunction affects a subset of Golgi enzymes and results in O-glycosylation defects

Each Golgi cisterna houses a specific set of different Golgi enzymes, ion channels, pH sensors, and transporters [62] [63] [64] [65] [66]. The Golgi enzymes catalyze the addition or removal of sugars to/from cargo glycoproteins and the addition of sulfate and phosphate groups [67]. Our previous study revealed that several tested Golgi enzymes, including B4GALT1, MGAT1, and GALNT2 were significantly depleted in GARP-KO cells [36]. Since these enzymes localize in different Golgi sub-compartments, we aim to determine if the decrease in their expression is a primary or secondary defect associated with VPS54 depletion. We observed that the reduction in protein level of B4GALT1 occurs only after a prolonged VPS54 depletion, indicating that this is not the immediate effect of GARP dysfunction (Figure 5A). Likewise, we observed no change in Golgi localization of B4GALT1 in cells acutely depleted for VPS54 (Figure 5B-C). Similar results were obtained with MGAT1 and GALNT2 (Figure S5A-D), suggesting that reduced protein stability of Golgi enzymes is an indirect consequence of GARP depletion. However, GARP acute depletion affected B4GALT1 localization to some extent since the localization of this enzyme became more sensitive to changes in Golgi pH induced by the chloroquine treatment (Figure S6), indicating that GARP activity is needed for proper *trans*-Golgi homeostasis, maybe via GARP-dependent stability of pH regulators such as TMEM87A.

**Figure 5:**
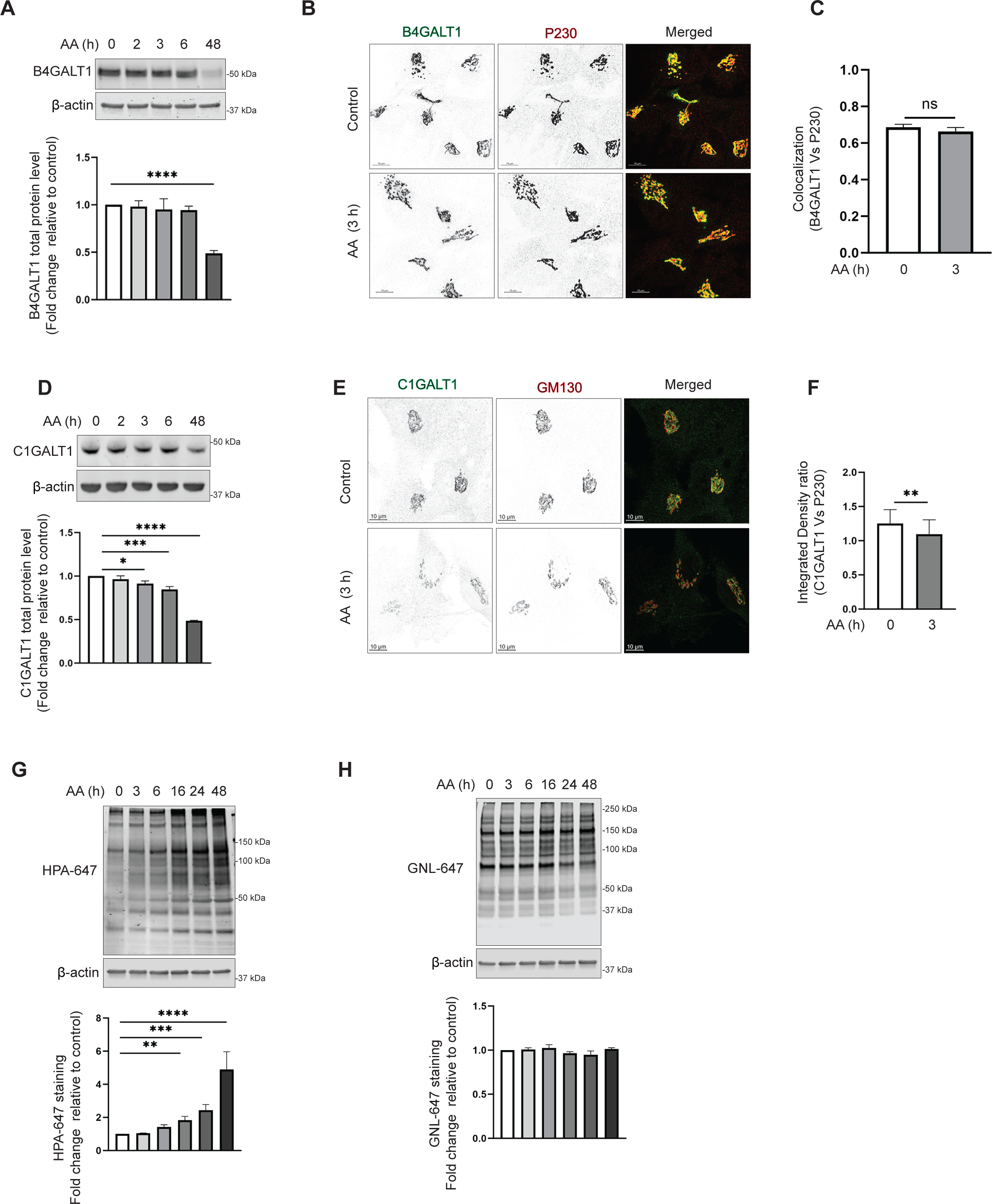
Acute VPS54 depletion affects a subset of Golgi enzymes and results in O-glycosylation defects. (A) WB analysis of cell lysates of AA treated RPE1 VPS54-mAID cells probed with (top panels) anti-B4GALT1 (A), and anti-C1GALT1 (D). β-actin was used as a loading control. The bottom panels on (A), and (D) are the quantification of the blots from three independent experiments. (B) Airyscan microscopy of RPE1 VPS54-mAID cells untreated (control) or treated with AA for 3 h and co-stained for B4GALT1 and P230. (C) Colocalization of B4GALT1 with P230 was determined by calculation of the Pearson’s correlation coefficient. (E) Airyscan microscopy of RPE1 VPS54-mAID cells untreated (control) or treated with AA for 3 h and co-stained for C1GALT1 and P230. (F) Integrated density ratio of C1GALT1 to P230 in control and AA treated group was determined using ImageJ. Statistical significance was calculated using paired t-test. ** p≤ 0.01. (G)Total proteins from AA treated RPE1 VPS54-mAID were resolved by SDS-PAGE and probed with (Top panel) HPA-647 (G), and GNL-647 (H). The bottom panels on (G), and (H) are the quantification of the blots from three independent experiments. Statistical significance was calculated using one-way ANOVA. ** p≤ 0.01, *** p≤ 0.001, **** p≤ 0.0001.

In agreement with the proposed GARP-related trans-Golgi dysfunction, we observed a significant decrease in the protein abundance of another enzyme, C1GALT1, within 3 h of the induction of VPS54 degradation (Figure 5D). Consistent with this, there was a decrease in colocalization of C1GALT1 with GM130 (Figure 5E-F). We reasoned that C1GALT1 mislocalization/degradation could lead to a specific defect in O-glycosylation. C1GALT1 transfers galactose from UDP-galactose to Tn antigen (GalNAcα1-O-Ser/Thr) to form core 1 *O*-glycan structure, T antigen. This step is critical for the biosynthesis of complex *O*-glycans [68]. *Helix pomatia* agglutin (HPA) binds to Tn antigen. Testing total cellular lysates in cells acutely depleted for VPS54 with HPA-647 lectin detected a significant increase in HPA-647 binding to several protein bands as early as 6 h after the induction of VPS54 degradation (Figure 5G), confirming C1GALT1 partial dysfunction. *O*-glycosylation abnormalities progressively increased upon a prolonged (16-48 hours) depletion of VPS54. The GARP-associated O-glycosylation defect appeared to be specific, as GNL-647 blot analysis did not reveal any abnormalities in proteins extracted from VPS54 acutely depleted cells, even after prolonged AA treatment (Figure 5H). This indicates that N-glycosylation defects are not a primary consequence of GARP dysfunction.

### GS15 is the Golgi SNARE that depends on GARP activity

SNAREs promote the fusion of vesicles containing cargo to their target membrane compartment. Once the TGN-derived transport vesicles are fused to the endosomal compartment, the SNAREs must return to the TGN as a normal process of recycling. Qc SNARE GS15/BET1L is shown to have increasing concentrations across the cisternae toward the *trans*-Golgi [69]. GS15 is believed to cycle via the endosomes, as it was found to be trapped in endosomes when endosome to Golgi recycling is disrupted [70]. In our study of VPS54-KO cells, we observed a significant decrease in total protein level and Golgi localization of GS15 [36]. We wondered if GS15 is sensitive to the rapid VPS54 degradation. Indeed, after 3 hours of VPS54 degradation induction, we observed that GS15 is mislocalized from the Golgi (Figure 6B-C). GS15 mislocalization led to significant depletion of GS15 protein (Figure 6A), indicating that Qc SNARE mislocalization and consequent degradation is a primary defect of GARP dysfunction. GS28/GOSR1 is a partner of GS15 in the STX5/GOSR1/BET1L/YKT6 SNARE complex and GS28 depletion led to GS15 instability [71]. Interestingly, we observed that GS28 protein stability or localization was not significantly altered upon VPS54 acute depletion, and its expression was decreased only after 48 h of GARP malfunction (Figure 6D and data not shown), indicating that GS15 relies on GARP function independently of its SNARE partner. Moreover, the stability of another GS15 SNARE partner, Qa SNARE STX5, was insensitive to GARP dysfunction (Figure 6F). GARP was shown to regulate the formation or stability of TGN STX16/STX6/VTI1A/VAMP4 SNARE complex [22]. Surprisingly, we found that the stability of STX6, VTI1A, VAMP4, and STX10 remained unaffected by VPS54 degradation (Figure 6F). These results indicate that GS15 is a unique Golgi SNARE protein, that relies on GARP for its localization and stability.

**Figure 6:**
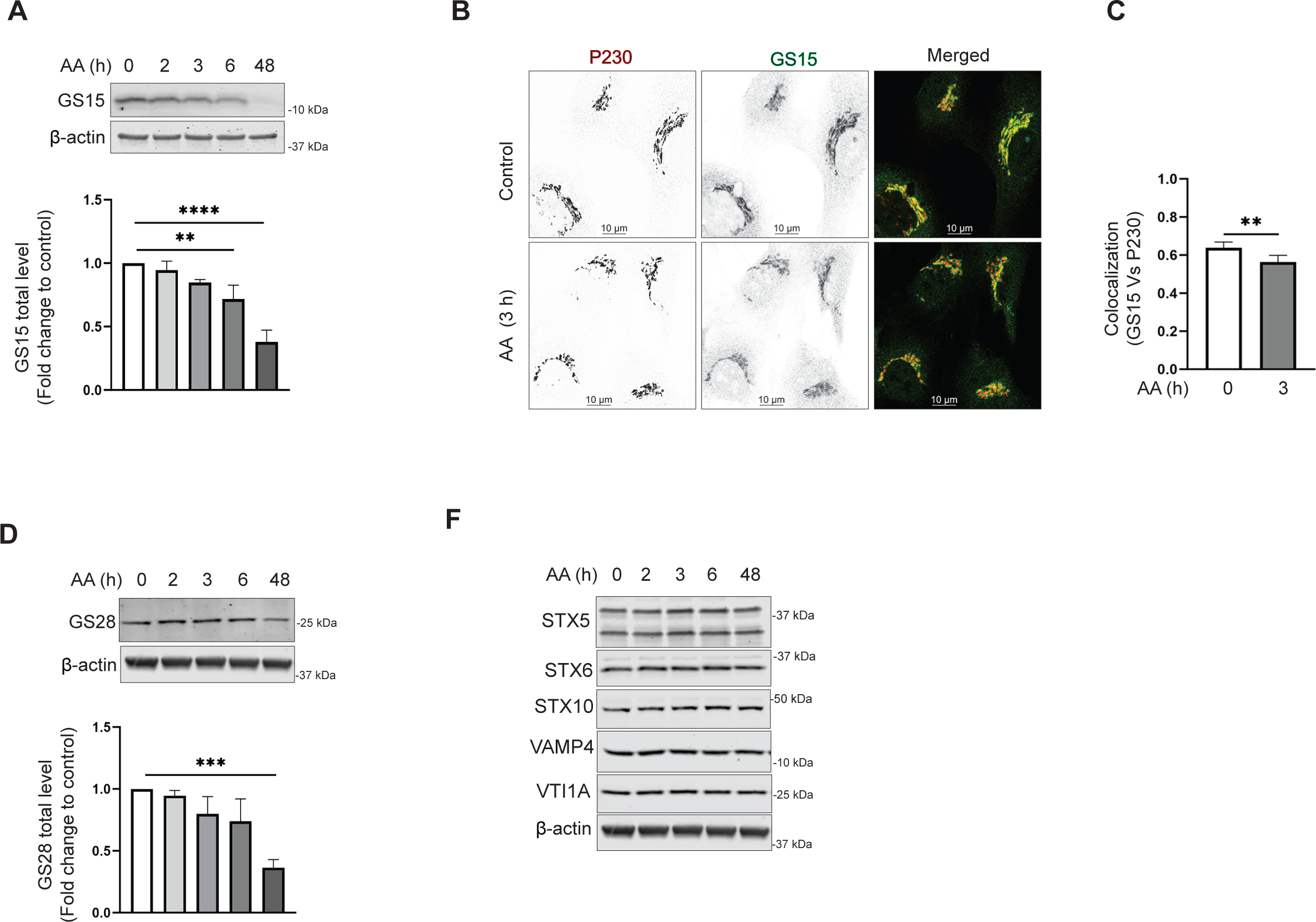
v-SNARE GS15 is mislocalized in VPS54-depleted cells. (A) RPE1 VPS54-mAID cells were treated with AA as indicated and cell lysates were probed with (top panel) anti-GS15 (A), and anti-GS28 (D). β-actin was used as a loading control. The bottom panels on (A), and (D) are the quantification of the blots from three independent experiments. (B) Airyscan microscopy of RPE1 VPS54-mAID cells untreated or treated with AA for 3 h and co-stained for GS15 and P230. (C) Colocalization analysis of GS15 and P230 was done by calculation of the Pearson’s correlation coefficient. Statistical significance was calculated using paired t-test. ** p≤ 0.01. (F) WB analysis of RPE1 VPS54-mAID cells treated with AA and probed with antibodies to STX5, STX6, STX10, VAMP4, and VTI1A respectively. β-actin was used as a loading control.

### Acute VPS54 depletion mislocalizes vesicular adaptor proteins and COPI coats

Previous investigation of GARP-KO cells revealed that several Golgi-located vesicular coats, including AP1, GGA, and COPI, were mislocalized to the cytosol and peripheral membranes. Coat binding to the Golgi membrane requires activation of ARF GTPases, facilitated by ARFGEF proteins GBF1, BIG1/ARFGEF1, and BIG2/ARFGEF2 [72]. In GARP-KO cells, ARFGEFs were mislocated from the Golgi ribbon to the cytosol and endolysosomal compartment [46]. To test if the Golgi coat localization defect is a primary manifestation of GARP malfunction, localization of β-adaptin (Figure 7A), GGA2 (Figure 7B), COPB2 (Figure 7C), and GBF1 (Figure 7D) was determined in cells acutely depleted for VPS54. Co-localization analysis revealed that 3 h of VPS54 depletion was sufficient for significant alterations in localization of all three coats and GBF1. In contrast, the localization of BIG1 was unchanged (Figure S7), suggesting that GBF1 mislocalization could be a primary defect that led to the malfunction of Golgi vesicular coats.

**Figure 7:**
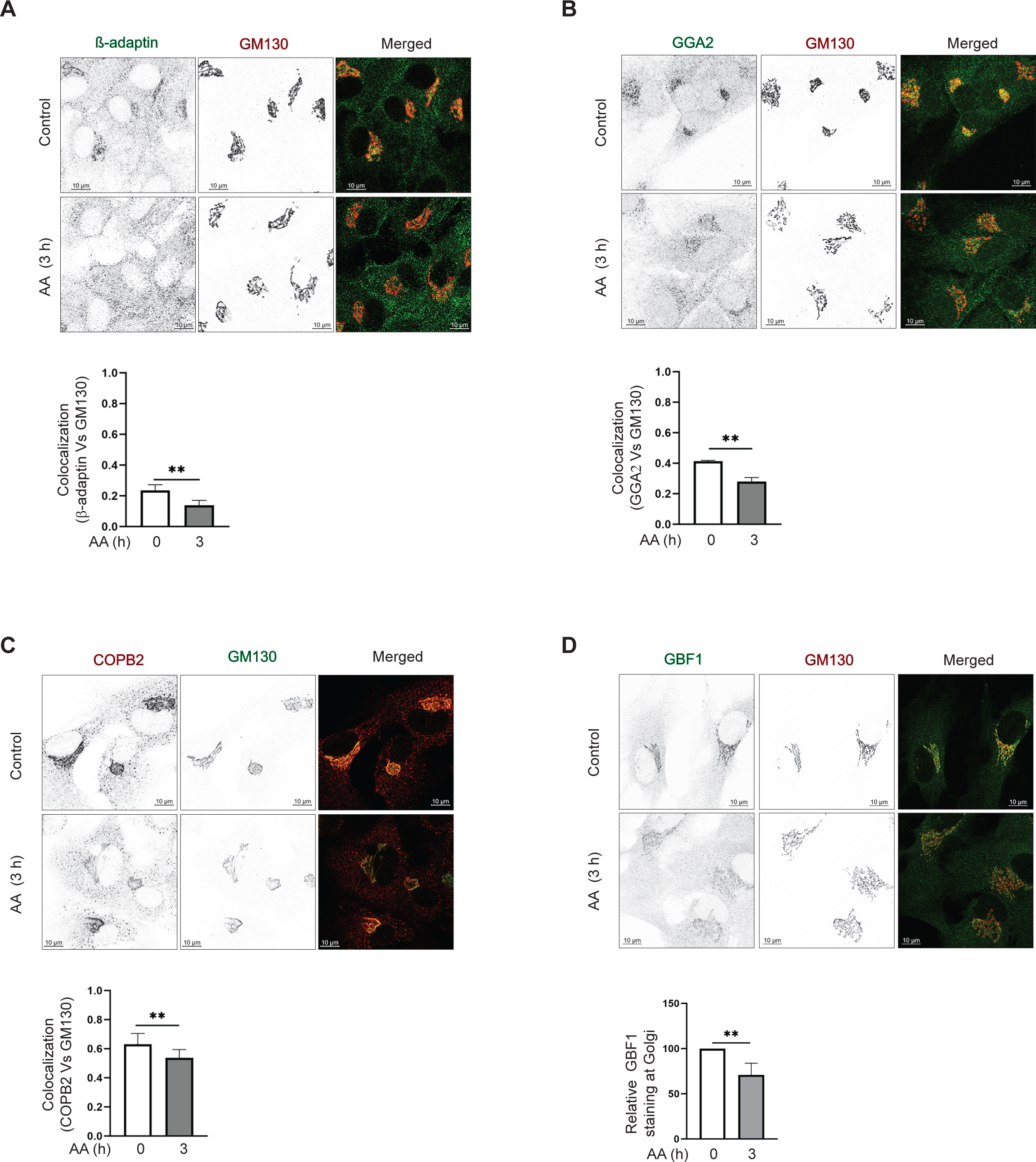
Acute VPS54 depletion mislocalizes vesicular adaptor proteins and COPI coats. (Top panel) Airyscan microscopy of RPE1 VPS54-mAID cells untreated and treated with AA for 3 h and co-stained for (A) β-adaptin and GM130 (B) GGA2 and GM130 (C) COPB2 and GM130 (D) GBF1 and GM130. (Bottom panel) Colocalization analysis of (A) β-adaptin and GM130. (B) GGA2 and GM130 (C) COPB2 and GM130 (D) GBF1 and GM130 was done by calculation of the Pearson’s correlation coefficient. ** p≤ 0.01.

### Rapid VPS54 depletion causes accumulation of GARP-dependent vesicles

Previous electron microscopy analysis of VPS54-KO cells revealed significant structural alterations in the Golgi complex, including swollen and partially fragmented cisternae. Notably, there was no substantial accumulation of small trafficking intermediates in GARP-KO cells, raising questions about GARP’s role as a vesicle tether [46]. To further investigate, we employed high-pressure freezing (HPF) and freeze substitution (FS) sample preparation for transmission electron microscopy (TEM) to identify early morphological changes in VPS54-depleted RPE1 cells. Our analysis revealed a significant increase in small vesicle-like structures (50-60 nm in diameter) near the Golgi (Figure 8A, B), supporting the role of GARP in vesicle tethering and suggesting that vesicle accumulation is an acute but transient phenotype of GARP complex dysfunction. Stalled GARP-dependent vesicles are likely to be cleared by autophagy, since we observed a number of autophagosomes in the Golgi area of VPS54-depleted cells. Some of these autophagosomes were filled with vesicle-like structures (Figure 8A, right panel shown by asterisk). Additionally, analysis of TEM images revealed the presence of large, round structures (0.2–0.6 microns in diameter) in the Golgi area. Electron-dense material frequently accumulates on one side of this organelle, possibly indicating the aggregation of lumenal cargo. The remainder of the Golgi stack appeared intact and not fragmented, indicating that the swollen Golgi in GARP-KO cells is likely a secondary manifestation of GARP dysfunction (Figure 8A). We hypothesized that the round structures represent altered TGN or enlarged late endosomal compartments resulting from depletion of components of the endosome-to-TGN recycling machinery. If this is the case, the enlarged structures must carry endosome-to-TGN receptors, like MPRs, which are known to recycle through this pathway [11].

**Figure 8:**
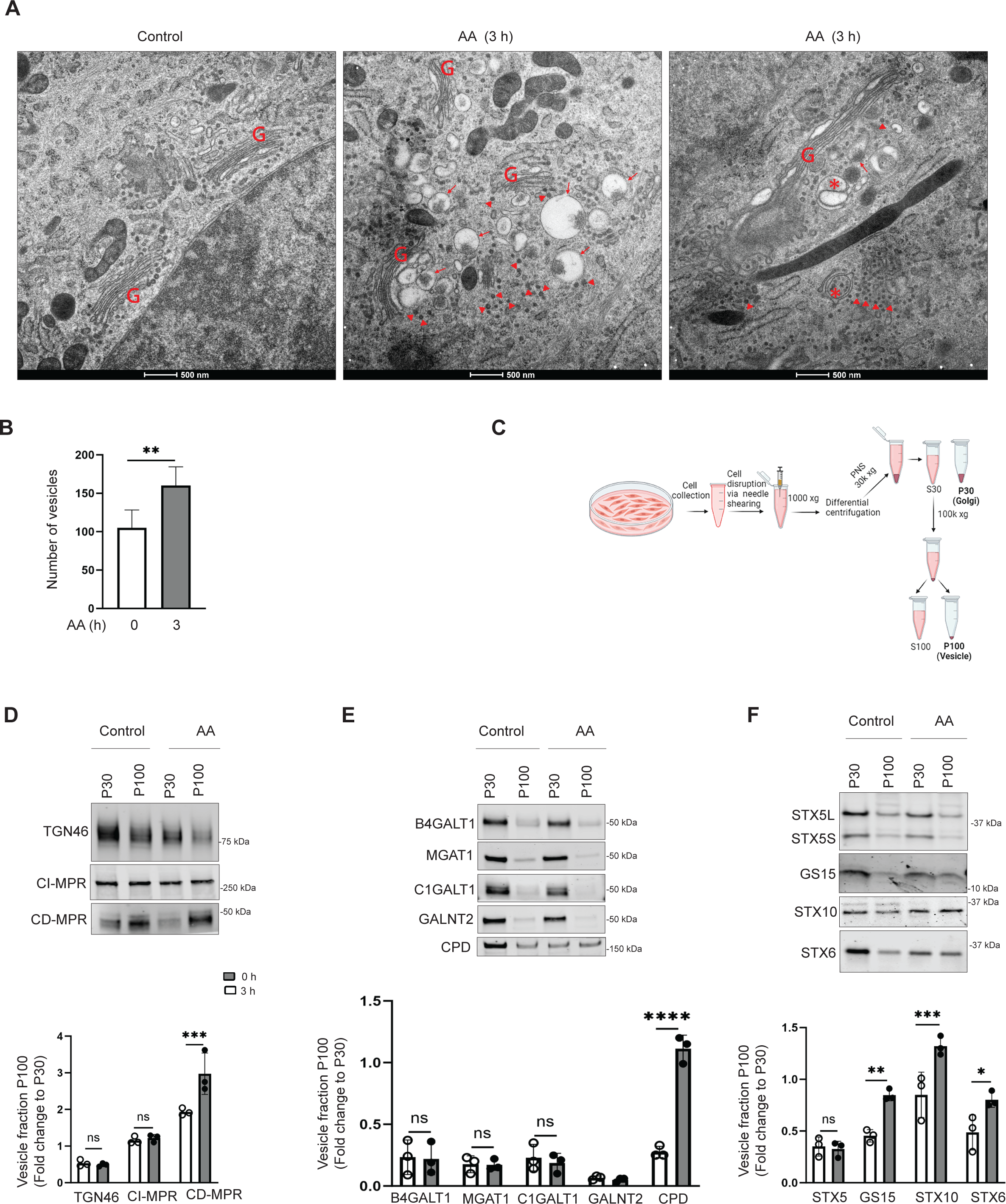
Rapid VPS54 depletion results in accumulation of GARP-dependent vesicles and alteration of TGN morphology. (A) Transmission Electron Microscopy of high-pressure frozen RPE1 VPS54-mAID cells grown on sapphire discs before and after 3 h of AA treatment. “G” indicates Golgi stacks. Arrowheads pointed to vesicle-like structures. Arrows indicate the enlarged vacuolar structures accumulated near the Golgi. Asterisks indicate the autophagosomes. Scale bar, 500 nm. (B) The graph represents the quantification of total number of vesicles around the Golgi before and after 3 h of AA treatment. (C) Schematic of cellular fractionation experiment to prepare P30 (Golgi), and P100 (Vesicle) fractions from control and AA treated groups. (D) WB analysis of TGN localized proteins (TGN46, CI-MPR, and CD-MPR) in Golgi and vesicle fractions. (E) WB analysis of Golgi enzymes (B4GALT1, MGAT1, C1GALT1, GALNT2, and CPD) in Golgi and vesicle fractions. (E) WB analysis of SNAREs (STX5, GS15, STX10, STX6) in Golgi and vesicle fractions.

To investigate whether GARP dysfunction leads to the accumulation of MPR in enlarged structures, we used VPS54-mAID cells stably expressing MPR-NeonGreen to do the live-cell imaging in control (Movie 1) and VPS54-depleted cells (Movie 2). Live-cell imaging of VPS54-depleted cells revealed the presence of MPR-NeonGreen signal in Golgi membranes, small vesicles, and large round organelles, similar to round structures observed by EM (Movie 2). We concluded that the accumulation of enlarged structures and small vesicles are primary defects associated with GARP dysfunction. To better understand the nature of GARP-dependent vesicles, RPE1 VPS54-mAID cells were mechanically disrupted and fractionated through differential centrifugation to separate Golgi membranes (P30) and vesicles (P100) (Figure 8C). We analyzed the distribution of three categories of proteins: endosome-TGN cycling receptors, Golgi enzymes, and Golgi SNAREs. Western blot analysis revealed that acute GARP inactivation led to the redistribution into the vesicular fraction of several proteins in these three categories, such as the recycling receptor CD-MPR (Figure 8D), the Golgi enzyme CPD (Figure 8E), and the Golgi SNAREs GS15 (Figure 8F), STX10 (Figure 8F), and STX6 (Figure 8F). Notably, the v-SNARE GS15 showed a significant increase in the vesicular pool following rapid GARP depletion (Figure 8F; Figure S8A), prompting us to use it to isolate the vesicles containing GS15 by native immunoprecipitation (Figure S8B). Western blot analysis demonstrated a 2.5-fold increase in GS15 protein in pull-down vesicles after acute GARP depletion (Figure S8C), while TGN46 levels decreased significantly (Figure S8D). A significant decrease in TGN46 signal in GS15 vesicles isolated from GARP-depleted cells likely indicates that the recycling pathways of TGN46 and GS15 are differently affected by VPS54 depletion (Figure S8D).

WB analysis of several Golgi enzymes, including B4GALT1, MGAT1, and GALNT2, did not reveal any significant changes in their abundance in GS15-positive vesicles isolated before and after VPS54 acute depletion (data not shown), but another Golgi enzyme, C1GALT1 was notably enriched in GARP-dependent vesicles, suggesting that its mislocalization contributes to the *O*-glycosylation defects observed in VPS54-depleted cells (Figure S8E). Additionally, the endosome-TGN SNAREs STX6 (Figure S8F) also showed a significant increase in GARP-dependent GS15-positive vesicles, despite no overall change in its total protein levels (Figure S8F).

Overall, the analysis of human cells acutely depleted for VPS54 revealed a marked increase in GS15-positive vesicles containing a subset of Golgi recycling proteins, highlighting a specific role for the GARP complex in the Golgi-endosomal trafficking cycle.

## Discussion

In this study, we have uncovered the immediate defects associated with GARP dysfunction and therefore distinguished between the primary and secondary defects, which are observed in previous studies of GARP knock-out and knock-down in mammalian cells [36] [46] [3] [11] [22]. We discovered that the mislocalization of vesicle coat proteins, increased number of GARP-dependent vesicles, alteration of *trans*-Golgi morphology, decreased stability and mislocalization of a endosome-to-TGN cycling proteins, and *O*-glycosylation as primary defects of GARP dysfunction (Figure 9).

**Figure 9:**
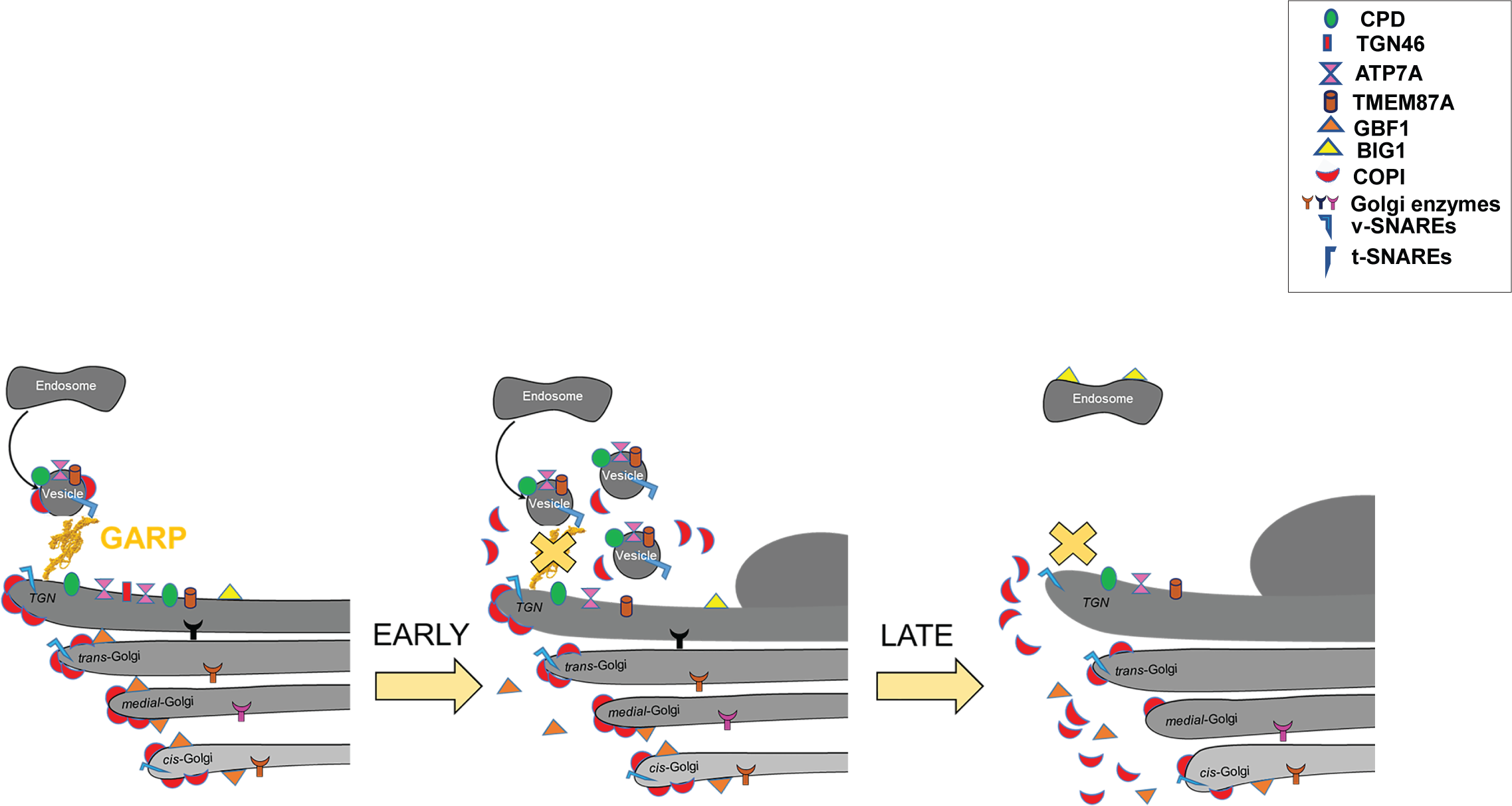
Cartoon depicting early and late defects associated with GARP dysfunction. (Left) Control cells with normal Golgi morphology, Golgi enzymes, endosome-Golgi trafficking proteins, and component of trafficking machineries such as coats and SNAREs. (Middle) 3 h post-depletion of VPS54, the trans-side of Golgi enlarged, vesicle number is increased, endosome-Golgi trafficking proteins are mislocalized and redistributed to vesicles, Golgi-associated coat proteins, and SNAREs are decreased. (Right) Severe alteration of Golgi structure, depletion of Golgi enzymes, coat proteins, and SNAREs are a late response.

While the degron activation resulted in a rapid depletion of VPS54, it did not change the total protein abundance of other subunits of GARP, supporting the notion that the EARP complex, which shares VPS51, VPS52, and VPS53 with GARP, is significantly more abundant than GARP [73]. In the future, it will be important to construct and test cells acutely depleted of both shared and unique EARP subunits to determine the specific roles of each complex in the TGN/endolysosomal trafficking cycle.

Mislocalization of three distinct protein coat complexes (COPI, AP-1, and GGA) was observed as one of the earliest phenotypes following VPS54 depletion. The dissociation of these coat complexes from the Golgi membrane is likely due to a reduction in ARF1-GTP levels, which are essential for the membrane association of all three vesicular coats [74]. This decrease in ARF1-GTP is likely caused by the mislocalization of the ARFGEF protein GBF1. It is intriguing that a malfunction in a TGN-localized trafficking factor primarily affects *cis-medial* GBF1 before influencing the *trans*-Golgi ARFGEF BIG1. GARP has been shown to be critical for cellular sphingolipid homeostasis [75], which may, in turn, influence phosphoinositide turnover through lipid exchange mechanisms at endoplasmic reticulum/TGN contact sites [76]. Since GBF1 binds to phosphoinositides, particularly PI3P, PI4P, and PI(4,5)P2, for membrane association [77], one possibility is that GARP depletion acutely disrupts the balance of Golgi phosphoinositides, thereby affecting GBF1 membrane binding. Another potential explanation for coat mislocalization could be the transient accumulation of non-tethered GARP-dependent vesicles.

Previous analysis of HeLa cells detected accumulation of CI-MPR in vesicle-like cytoplasmic staining and in a light membrane fraction in VPS52-RNAi-depleted cells [11], suggesting the buildup of small trafficking carriers. However, more recent EM studies of GARP knockout cells did not detect vesicle accumulation [46], raising questions about the role of GARP in vesicle tethering. Both RNAi and CRISPR/Cas9 KO techniques require several days for protein depletion, which could introduce artifacts and/or allow cellular adaptation to the loss of the target protein. The detection of GARP-dependent vesicles by EM in cells acutely depleted for VSP54, along with the identification of specific cargo proteins associated with these trafficking carriers, provides the first direct experimental evidence supporting GARP’s role as a vesicular tether. EM data also suggested that accumulated GARP-dependent vesicles are getting removed by autophagy, resolving the discrepancy between phenotypes in acutely depleted versus GARP-KO cells.

What is the protein cargo of GARP-dependent transport carriers? Our analysis identified several transmembrane Golgi resident proteins whose abundance and/or localization were significantly affected by acute VPS54 depletion. The list includes TGN46, ATP7A, TMEM87A, CPD, CD-MPR, C1GALT1, GS15, STX6 and STX10. Consistent with previous reports in VPS54-KO cells, the TGN46 protein was highly sensitive to VPS54 depletion. We found that TGN46 began to mislocalize from the Golgi into punctate structures within one hour of inducing VPS54 degradation (unpublished data), making TGN46 the fastest responder to VPS54 loss. This suggests that TGN46 may cycle between the TGN and endolysosomal compartments at a rapid rate, and GARP dysfunction quickly leads to its degradation in lysosomes. Supporting this hypothesis, pretreatment of GARP deficient cells with protease inhibitors partially rescued TGN46 expression. Although TGN46 was detected in immunoprecipitated GS15 vesicles, GARP malfunction did not lead to accumulation of TGN46 in GS15 carriers, indicating that the trafficking itinerary of this putative cargo receptor is distinct from that of GS15. Future investigations monitoring transport carriers via live microscopy using fluorescently tagged TGN46 in GARP-depleted cells should help clarify this issue.

ATP7A, also known as Menke’s protein, it’s predominantly localized at the TGN and it’s responsible for regulation of copper homeostasis in the cell [6] [78] [79] [80] [81]. In the steady state, ATP7A is in the Golgi, but when the cellular copper level is high, ATP7A migrates to the plasma membrane and regulates intracellular copper levels [82]. A study by Heather *et al*., showed that ATP7A interacts with the COG complex [83] and the ablation of the COG complex downregulated ATP7A in mammalian cells [84], indicating rapid intra-Golgi recycling of this copper transporter. ATP7A has also been shown to physically interact with AP-1 coat complex and that AP-1 regulates ATP7A localization under basal copper concentrations [85]. AP-1 is preferentially regulating endosome to Golgi retrograde trafficking [86] and it is likely that GARP depletion blocks constitutive endosome/Golgi recycling of ATP7A leading to its mistargeting and degradation. Another GARP sensitive protein, TMEM87A, which appears to play a critical role in maintaining Golgi pH, and its knockout in mice leads to Golgi fragmentation and altered protein glycosylation [57] [87]. Furthermore, overexpression of TMEM87A in VPS54-KO cells partially restored retrograde transport from endosomes to the TGN [3]. Notably, GARP-deficient cells became hypersensitive to chloroquine treatment that elevated Golgi pH. The specific mislocalization of C1GALT1 enzyme, along with TMEM87A depletion-related changes in Golgi pH, may contribute to *O*-glycosylation defects in VPS54-depleted cells.

It remains unclear whether TGN golgins and GARP regulate distinct or overlapping trafficking pathways. On the one hand, golgin-decorated mitochondria can attract trafficking intermediates carrying CI-MPR [20], a receptor unaffected by acute VPS54 depletion, suggesting that GARP and golgins may be involved in tethering different membrane carriers. On the other hand, both GARP and TGN golgin membrane recruitment is regulated by the same small GTPase, ARFRP1 [31], and our preliminary data show a very close spatial proximity between golgins and GARP (data not shown), pointing to a possible coordination between the two. Future investigations into membrane trafficking in cells deficient in both golgins and GARP will help clarify whether there is redundancy within the TGN tethering machinery [88].

Our data indicated that acute depletion of VPS54 specifically affected the sorting of the lysosomal enzyme cathepsin D, resulting in increased secretion of its precursor. However, we observed that the localization of CI-MPR remained unchanged, while CD-MPR mostly stayed in the Golgi proper, challenging the notion that GARP is directly involved in the trafficking of MPR proteins in human cells. One explanation of the microscopy data is that the tethering of MPR-carrying intermediates is primarily mediated by TGN golgins [20], but biochemical data suggest a more complex scenario. Cellular fractionation of VPS54-depleted cells showed that some CD-MPR, but not CI-MPR, was redistributed to the GARP-dependent vesicle fraction, indicating that the two mannose-6-phosphate receptors may follow different TGN-endosome pathways. It is plausible that the partial mislocalization of CD-MPR, along with potential changes in TGN acidity due to the mislocalization of TMEM87A, could be a primary cause of the missorting of pro-cathepsin D.

Interestingly, we found VPS54 acute depletion also affected the secretion of fibronectin (FN1). The increase in secretion of FN1 was not due to the increase in its expression. One of the reasons for the increase secretion of FN1 could be related to the altered pH and/or morphology of the TGN. A study by Mayuko *et al.* showed that cells with altered Golgi morphology stimulate the transport of secretory alkaline phosphatase [89], suggesting the importance of Golgi morphology and GARP machinery in controlling the rate and quality of protein secretion. This potential GARP function is in agreement with the recently discovered AP-1 driven cycling of secretory cargo in yeast cells [90].

How many types of trafficking intermediates are regulated by GARP complex? Initial characterization of GS15-containing vesicles revealed accumulation of three GARP-dependent proteins, Golgi enzyme C1GALT1 and two SNAREs GS15 and STX6. However, the abundance of TGN46 was significantly decreased in GS15-containing carriers isolated from VPS54 deprived cells, indicating that at least some of TGN46 is returned from post-Golgi compartments by different carriers.

The exact composition and distribution of vesicle fusion machinery regulated by GARP is another question. Qc SNARE GS15 is accumulated in GARP-dependent trafficking intermediates, but GS15-KO results in phenotypes much milder than VPS54-KO [46] [71] and GARP is not known to regulate SNARE GS15-containing complexes. Instead, GARP is predicted to regulate STX16/STX6/VTI1A/VAMP4 SNARE complex [22], but, intriguingly, STX16 SNARE assembly is also regulated by COG vesicle tethering complex [91]. Moreover, cells acutely depleted of VPS54 did not show any mislocalization or degradation of R-SNARE VAMP4, while Qa–SNARE STX16 is accumulated in GS15-independent vesicle carriers (data not shown). Future proteomic analysis of GARP-dependent trafficking intermediates should clarify these important questions.

In summary, analysis of human cells acutely depleted for VPS54 revealed key cellular defects linked to GARP dysfunction (Table 3). Future proteomic studies on GARP-dependent trafficking intermediates, combined with proximity labeling and in vitro methods, will enhance our understanding of GARP’s role in membrane trafficking.

**Table 3.** Golgi proteins affected by acute and prolonged GARP depletion.

| Affected proteins | Early defects<br>(up to six hours of VPS54<br>depletion) | Late defects<br>(more than six hours of<br>VPS54 depletion or KO) |
| --- | --- | --- |
| TGN proteins | TGN46<br>ATP7A<br>TMEM87A<br>CPD |  |
| Golgi SNAREs | GS15/BET1L | GS28/GOSR1 |
| Golgi enzymes | C1GALT1 | B4GALT1<br>MGAT1<br>GALNT2 |
| Calcium homeostasis<br>maintenance proteins |  | SDF4<br>ATP2C1 |
| ARFGEFs | GBF1 | BIG1 |
| Coats/adaptors/accessory<br>proteins | β-adaptin<br>GGA2<br>COPB2 | |

## Acknowledgments

We are very immensely thankful to Juan S. Bonifacino for providing the HeLa VPS54-KO cell line and plasmids used in the study. We acknowledge Tetyana Kudlyk’s contribution to creating the cell lines and Farhana Taher Sumya for preparing lentivirus expressing OsTIR1 (F74G)-V5. We would like to be thankful to Eric Campeau, Wei Guo, Paul Kaufman, Frank Perez, Santiago M. Di Pietro, Didier Trono, and others who provided reagents and cell lines. We are grateful to all members of Lupashin’s lab and Roy Morello for their comments on the manuscript. This work was supported by the National Institute of Health (R01GM083144) and by UAMS Easy Win Early Victory grant program (VL).

## Supplementary Figure legends

**Figure S1: VPS54-mAID rescue Golgi localization defects in VPS54-KO HeLa cells.** HeLa cells expressing VPS54-mAID and VPS54-KO were stained with antibodies to TGN46 (A), GBF1 (B), and COPB2 (C) and analyzed by confocal microscopy.

**Figure S2: Efficient depletion of VPS54 in HeLa cells expressing VPS54-mAID following AA treatment.** (A) WB analysis of HeLa cells expressing VPS54-mAID following rapid VPS54 depletion for 0, 0.5, 1, 2, 3, 24, and 48 h respectively and probed with anti-myc (VPS54) antibody. The bottom panel shows the quantification of the blot. (B) HeLa cells expressing VPS54-mAID were co-stained with myc (red) and B4GALT1 (green) after 3 h of VPS54 depletion.

**Figure S3: Effect of VPS54-KO on ATP7A and CPD localization.** (A) (Top panel) WB analysis of RPE1 cells expressing VPS54-mAID before and after the treatment of AA and protease inhibitor (PI) and probed with anti-TGN46. (Bottom panel) Quantification of the blots from three independent experiments. Statistical significance was assessed using one-way ANOVA. **** p≤ 0.0001, ** p≤ 0.01. (B) RPE1 cells expressing VPS54-mAID were transiently transfected with LAMP2-mCherry and TGN38-GFP for 24 h followed by 3 h AA treatment and cells were imaged using airyscan microscopy. (C) Airyscan microscopy images of RPE1 WT, VPS54-KO, and VPS54-mAID cells stained with anti-B4GALT1, anti-GM130, and anti-ATP7A antibodies. (D) Co-staining of RPE1 WT and VPS54-KO cells for CPD and P230 using anti-CPD and anti-P230 antibodies.

**Figure S5: Acute VPS54 depletion does not affect the Golgi enzymes MGAT1 and GALNT2 in RPE1 cells.** (A) WB analysis of total lysates of RPE1cells expressing VPS54-mAID were treated with AA and probed with (Top panel) anti-MGAT1 (A), and anti-GALNT2 (D). β-actin was used as a loading control. The bottom panels on (A), and (D) are the quantification of the blots from three independent experiments. (B) Airyscan microscopy of RPE1 VPS54-mAID cells untreated or treated with AA for 3 h and co-stained for MGAT1 and P230. (C) Colocalization analysis of MGAT1 with P230 was done by calculation of the Pearson’s correlation coefficient.

**Figure S6: Rapid depletion of VPS54 stimulates B4GALT1 relocation to endosomes following CQ treatment.** (A) IF images of RPE1 cells expressing VPS54-mAID. Untreated cells (control), or treated with CQ for 3 hours (CQ), or treated with AA for 1 hour followed by CQ treatment for 3 hours (AA+CQ), or treated with AA and CQ and recovered for 3 hours (AA+CQ+W) or treated with CQ and recovered for 3 hours (CQ+W) were stained for GPP130, B4GALT1, and Giantin. (B) Colocalization analysis of B4GALT1 and Giantin was done by calculation of the Pearson’s correlation coefficient. Statistical significance was calculated using one-way ANOVA. ** p≤ 0.01, *** p≤ 0.001, **** p≤ 0.0001. (C) Colocalization analysis of GPP130 and Giantin was also done by calculation of the Pearson’s correlation coefficient. Statistical significance was calculated using one-way ANOVA.

**Figure S7: Acute VPS54 depletion does not affect BIG1 localization.** (A) Airyscan microscopy of RPE1 VPS54-mAID cells untreated or treated with AA for 3 h and co-stained for BIG1 and P230. (B) Colocalization of BIG1 and P230 was determined by the calculation of the Pearson’s correlation coefficient.

**Figure S8: Acute VPS54 depletion led to accumulation of C1GALT1 and STX6 in the GS15 positive vesicles.** (A) WB analysis of GS15 in Golgi and vesicle fraction before (-) and after (+) acute VPS54 depletion. (B) Schematic of GS15 IP from the vesicular fraction (S30) using GS15 antibody in control and AA treated groups. (C-F) (Top panel) WB analysis of GS15 IP in control and AA treated groups probed with anti-GS15 (C), TGN46 (D), C1GALT1 (E), and STX6 (F). (C-F) (Bottom panel) The graph represents the quantification of three independent blots.

**Movie 1: Super-resolution airyscan live cell imaging of RPE1 cells stably expressing CD-MPR-neon green**. RPE1 VPS54-mAID cells were stably expressed with CD-MPR-neon green and imaged in real-time in the absence of AA. Live cell imaging was done for 60 frames in 30 seconds.

**Movie 2: Super-resolution airyscan live cell imaging of RPE1 cells stably expressing CD-MPR-neon green treated with AA for 3 hours**. RPE1 VPS54-mAID cells were stably expressed with CD-MPR-neon green was treated with AA for 3 h and imaged in real-time (AA 3 h). Live cell imaging was done for 60 frames in 30 seconds. Arrows on VPS54 depleted cells showed the accumulation of CD-MPR in the enlarged round compartment near to the Golgi. The time stamp represents seconds.

## Notes

### Competing Interest Statement

The authors have declared no competing interest.

### Summary of Updates

This version of the manuscript has been modified to correct several grammatical errors and add Movie files missing in the first submission

## References

1. Reaves B, Horn M, Banting G (1993) TGN38/41 recycles between the cell surface and the TGN: brefeldin A affects its rate of return to the TGN. Molecular biology of the cell 4 (1):93–105

2. Ladinsky MS, Howell KE (1992) The trans-Golgi network can be dissected structurally and functionally from the cisternae of the Golgi complex by brefeldin A. European journal of cell biology 59 (1):92–105

3. Hirata T, Fujita M, Nakamura S, Gotoh K, Motooka D, Murakami Y, Maeda Y, Kinoshita T (2015) Post-Golgi anterograde transport requires GARP-dependent endosome-to-TGN retrograde transport. Molecular biology of the cell 26 (17):3071–3084

4. Tu Y, Zhao L, Billadeau DD, Jia D (2020) Endosome-to-TGN trafficking: organelle-vesicle and organelle-organelle interactions. Frontiers in cell and developmental biology 8:163

5. Petris M, Mercer J, Culvenor J, Lockhart P, Gleeson P, Camakaris J (1996) Ligand-regulated transport of the Menkes copper P-type ATPase efflux pump from the Golgi apparatus to the plasma membrane: a novel mechanism of regulated trafficking. The EMBO journal 15 (22):6084–6095

6. Zhu S, Shanbhag V, Hodgkinson VL, Petris MJ (2016) Multiple di-leucines in the ATP7A copper transporter are required for retrograde trafficking to the trans-Golgi network. Metallomics 8 (9):993–1001

7. Ruturaj R, Mishra M, Saha S, Maji S, Rodriguez-Boulan E, Schreiner R, Gupta A (2024) Regulation of the apico-basolateral trafficking polarity of the homologous copper-ATPases ATP7A and ATP7B. Journal of Cell Science 137 (5)

8. Varlamov O, Fricker LD (1998) Intracellular trafficking of metallocarboxypeptidase D in AtT-20 cells: localization to the trans-Golgi network and recycling from the cell surface. Journal of Cell Science 111 (7):877–885

9. Cattin-Ortolá J, Kaufman JG, Gillingham AK, Wagstaff JL, Peak-Chew S-Y, Stevens TJ, Boulanger J, Owen DJ, Munro S (2024) Cargo selective vesicle tethering: The structural basis for binding of specific cargo proteins by the Golgi tether component TBC1D23. Science Advances 10 (13):eadl0608

10. Chia PZC, Gasnereau I, Lieu ZZ, Gleeson PA (2011) Rab9-dependent retrograde transport and endosomal sorting of the endopeptidase furin. Journal of Cell Science 124 (14):2401–2413

11. Pérez-Victoria FJ, Mardones GA, Bonifacino JS (2008) Requirement of the human GARP complex for mannose 6-phosphate-receptor-dependent sorting of cathepsin D to lysosomes. Molecular biology of the cell 19 (6):2350–2362

12. Pan X, Zaarur N, Singh M, Morin P, Kandror KV (2017) Sortilin and retromer mediate retrograde transport of Glut4 in 3T3-L1 adipocytes. Molecular biology of the cell 28 (12):1667–1675

13. Dumanis SB, Burgert T, Caglayan S, Füchtbauer A, Füchtbauer E-M, Schmidt V, Willnow TE (2015) Distinct functions for anterograde and retrograde sorting of SORLA in amyloidogenic processes in the brain. Journal of Neuroscience 35 (37):12703–12713

14. Matsudaira T, Niki T, Taguchi T, Arai H (2015) Transport of the cholera toxin B-subunit from recycling endosomes to the Golgi requires clathrin and AP-1. Journal of Cell Science 128 (16):3131–3142

15. Mallard F, Antony C, Tenza D, Salamero J, Goud B, Johannes L (1998) Direct pathway from early/recycling endosomes to the Golgi apparatus revealed through the study of shiga toxin B-fragment transport. The Journal of cell biology 143 (4):973–990

16. Smith RD, Willett R, Kudlyk T, Pokrovskaya I, Paton AW, Paton JC, Lupashin VV (2009) The COG complex, Rab6 and COPI define a novel Golgi retrograde trafficking pathway that is exploited by SubAB toxin. Traffic 10 (10):1502-1517

17. Bonifacino JS, Glick BS (2004) The mechanisms of vesicle budding and fusion. cell 116 (2):153-166

18. Whyte JR, Munro S (2002) Vesicle tethering complexes in membrane traffic. Journal of Cell Science 115 (13):2627–2637

19. Rothman JE (1996) The protein machinery of vesicle budding and fusion. Protein science 5 (2):185–194

20. Wong M, Munro S (2014) The specificity of vesicle traffic to the Golgi is encoded in the golgin coiled-coil proteins. Science 346 (6209):1256898

21. Cheung P-yP, Pfeffer SR (2016) Transport vesicle tethering at the trans Golgi network: coiled coil proteins in action. Frontiers in cell and developmental biology 4:18

22. Pérez-Victoria FJ, Bonifacino JS (2009) Dual roles of the mammalian GARP complex in tethering and SNARE complex assembly at the trans-golgi network. Molecular and cellular biology.

23. Liewen H, Meinhold-Heerlein I, Oliveira V, Schwarzenbacher R, Luo G, Wadle A, Jung M, Pfreundschuh M, Stenner-Liewen F (2005) Characterization of the human GARP (Golgi associated retrograde protein) complex. Experimental cell research 306 (1):24–34

24. Koumandou VL, Dacks JB, Coulson RM, Field MC (2007) Control systems for membrane fusion in the ancestral eukaryote; evolution of tethering complexes and SM proteins. BMC evolutionary biology 7:1–17

25. Oka T, Krieger M (2005) Multi-component protein complexes and Golgi membrane trafficking. Journal of biochemistry 137 (2):109–114

26. Bröcker C, Engelbrecht-Vandré S, Ungermann C (2010) Multisubunit tethering complexes and their role in membrane fusion. Current Biology 20 (21):R943–R952

27. Santana-Molina C, Gutierrez F, Devos DP (2021) Homology and modular evolution of CATCHR at the origin of the eukaryotic endomembrane system. Genome Biology and Evolution 13 (7):evab125

28. Conibear E, Stevens TH (2000) Vps52p, Vps53p, and Vps54p form a novel multisubunit complex required for protein sorting at the yeast late Golgi. Molecular biology of the cell 11 (1):305-323

29. Siniossoglou S, Pelham HR (2001) An effector of Ypt6p binds the SNARE Tlg1p and mediates selective fusion of vesicles with late Golgi membranes. The EMBO journal

30. Gershlick DC, Ishida M, Jones JR, Bellomo A, Bonifacino JS, Everman DB (2019) A neurodevelopmental disorder caused by mutations in the VPS51 subunit of the GARP and EARP complexes. Human molecular genetics 28 (9):1548–1560

31. Ishida M, Bonifacino JS (2019) ARFRP1 functions upstream of ARL1 and ARL5 to coordinate recruitment of distinct tethering factors to the trans-Golgi network. The Journal of cell biology 218 (11):3681

32. Khakurel A, Lupashin VV (2023) Role of GARP vesicle tethering complex in Golgi physiology. International journal of molecular sciences 24 (7):6069

33. Gershlick DC, Schindler C, Chen Y, Bonifacino JS (2016) TSSC1 is novel component of the endosomal retrieval machinery. Molecular biology of the cell 27 (18):2867–2878

34. Abascal-Palacios G, Schindler C, Rojas AL, Bonifacino JS, Hierro A (2013) Structural basis for the interaction of the Golgi-Associated Retrograde Protein Complex with the t-SNARE Syntaxin 6. Structure 21 (9):1698–1706

35. Khakurel A, Kudlyk T, Lupashin VV (2022) Generation and analysis of hTERT-RPE1 VPS54 knock-out and rescued cell lines. In: Golgi: Methods and Protocols. Springer, pp 349–364

36. Khakurel A, Kudlyk T, Bonifacino JS, Lupashin VV (2021) The Golgi-associated retrograde protein (GARP) complex plays an essential role in the maintenance of the Golgi glycosylation machinery. Molecular biology of the cell 32 (17):1594–1610

37. Yesbolatova A, Saito Y, Kitamoto N, Makino-Itou H, Ajima R, Nakano R, Nakaoka H, Fukui K, Gamo K, Tominari Y (2020) The auxin-inducible degron 2 technology provides sharp degradation control in yeast, mammalian cells, and mice. Nature communications 11 (1):5701

38. Saito Y, Kanemaki MT (2021) Targeted Protein Depletion Using the Auxin-Inducible Degron 2 (AID2) System. Current Protocols 1 (8):e219

39. Nishimura K, Fukagawa T, Takisawa H, Kakimoto T, Kanemaki M (2009) An auxin-based degron system for the rapid depletion of proteins in nonplant cells. Nature methods 6 (12):917–922

40. Holland AJ, Fachinetti D, Han JS, Cleveland DW (2012) Inducible, reversible system for the rapid and complete degradation of proteins in mammalian cells. Proceedings of the National Academy of Sciences 109 (49):E3350–E3357

41. Ambrosio AL, Boyle JA, Di Pietro SM (2012) Mechanism of platelet dense granule biogenesis: study of cargo transport and function of Rab32 and Rab38 in a model system. Blood, The Journal of the American Society of Hematology 120 (19):4072–4081

42. Sanjana NE, Shalem O, Zhang F (2014) Improved vectors and genome-wide libraries for CRISPR screening. Nature methods 11 (8):783–784

43. Natsume T, Kiyomitsu T, Saga Y, Kanemaki MT (2016) Rapid protein depletion in human cells by auxin-inducible degron tagging with short homology donors. Cell Reports 15 (1):210–218

44. Campeau E, Ruhl VE, Rodier F, Smith CL, Rahmberg BL, Fuss JO, Campisi J, Yaswen P, Cooper PK, Kaufman PD (2009) A versatile viral system for expression and depletion of proteins in mammalian cells. PLoS One 4 (8):e6529

45. Dull T, Zufferey R, Kelly M, Mandel R, Nguyen M, Trono D, Naldini L (1998) A third-generation lentivirus vector with a conditional packaging system. Journal of virology 72 (11):8463–8471

46. Khakurel A, Kudlyk T, Pokrovskaya I, D’Souza Z, Lupashin VV (2022) GARP dysfunction results in COPI displacement, depletion of Golgi v-SNAREs and calcium homeostasis proteins. Frontiers in cell and developmental biology 10:1066504

47. Sumya FT, Pokrovskaya ID, Lupashin V (2021) Development and initial characterization of cellular models for COG complex-related CDG-II diseases. Frontiers in genetics 12:733048

48. Buser DP, Spang A (2023) Protein sorting from endosomes to the TGN. Frontiers in cell and developmental biology 11:1140605

49. Lujan P, Garcia-Cabau C, Wakana Y, Lillo JV, Rodilla-Ramírez C, Sugiura H, Malhotra V, Salvatella X, Garcia-Parajo MF, Campelo F (2024) Sorting of secretory proteins at the trans-Golgi network by human TGN46. Elife 12:RP91708

50. Banting G, Ponnambalam S (1997) TGN38 and its orthologues: roles in post-TGN vesicle formation and maintenance of TGN morphology. Biochimica et Biophysica Acta (BBA)-Molecular Cell Research 1355 (3):209–217

51. Bos K, Wraight C, Stanley K (1993) TGN38 is maintained in the trans-Golgi network by a tyrosine-containing motif in the cytoplasmic domain. The EMBO journal 12 (5):2219–2228

52. Mallet WG, Maxfield FR (1999) Chimeric forms of furin and TGN38 are transported from the plasma membrane to the trans-Golgi network via distinct endosomal pathways. The Journal of cell biology 146 (2):345–360

53. Humphrey JS, Peters PJ, Yuan LC, Bonifacino JS (1993) Localization of TGN38 to the trans-Golgi network: involvement of a cytoplasmic tyrosine-containing sequence. The Journal of cell biology 120 (5):1123–1135

54. Reaves B, Banting G (1994) Overexpression of TGN38/41 leads to mislocalisation of γ-adaptin. FEBS letters 351 (3):448–456

55. La Fontaine S, Mercer JF (2007) Trafficking of the copper-ATPases, ATP7A and ATP7B: role in copper homeostasis. Archives of biochemistry and biophysics 463 (2):149-167

56. Polishchuk R, Lutsenko S (2013) Golgi in copper homeostasis: a view from the membrane trafficking field. Histochemistry and cell biology 140:285–295

57. Kang H, Han A-R, Zhang A, Jeong H, Koh W, Lee JM, Lee H, Jo HY, Maria-Solano MA, Bhalla M (2024) GolpHCat (TMEM87A), a unique voltage-dependent cation channel in Golgi apparatus, contributes to Golgi-pH maintenance and hippocampus-dependent memory. Nature communications 15 (1):5830

58. Harasaki K, Lubben NB, Harbour M, Taylor MJ, Robinson MS (2005) Sorting of major cargo glycoproteins into clathrin-coated vesicles. Traffic 6 (11):1014–1026

59. Ghosh P, Dahms NM, Kornfeld S (2003) Mannose 6-phosphate receptors: new twists in the tale. Nature Reviews Molecular Cell Biology 4 (3):202–213

60. Olson LJ, Hindsgaul O, Dahms NM, Kim J-JP (2008) Structural insights into the mechanism of pH-dependent ligand binding and release by the cation-dependent mannose 6-phosphate receptor. Journal of Biological Chemistry 283 (15):10124–10134

61. Bohnsack RN, Song X, Olson LJ, Kudo M, Gotschall RR, Canfield WM, Cummings RD, Smith DF, Dahms NM (2009) Cation-independent Mannose 6-Phosphate Receptor. Journal of Biological Chemistry 284 (50):35215–35226

62. Lissandron V, Podini P, Pizzo P, Pozzan T (2010) Unique characteristics of Ca2+ homeostasis of the trans-Golgi compartment. Proceedings of the National Academy of Sciences 107 (20):9198–9203

63. Munro S (2005) The Golgi apparatus: defining the identity of Golgi membranes. Current opinion in cell biology 17 (4):395–401

64. Stanley P (2011) Golgi glycosylation. Cold Spring Harbor perspectives in biology 3 (4):a005199

65. Li J, Wang Y (2022) Golgi metal ion homeostasis in human health and diseases. Cells 11 (2):289

66. Kellokumpu S (2019) Golgi pH, ion and redox homeostasis: how much do they really matter? Frontiers in cell and developmental biology 7:93

67. Hirschberg CB, Robbins PW, Abeijon C (1998) Transporters of nucleotide sugars, ATP, and nucleotide sulfate in the endoplasmic reticulum and Golgi apparatus. Annual review of biochemistry 67 (1):49–69

68. Sun X, Zhan M, Sun X, Liu W, Meng X (2021) C1GALT1 in health and disease. Oncology letters 22 (2):1–15

69. Volchuk A, Ravazzola M, Perrelet A, Eng WS, Di Liberto M, Varlamov O, Fukasawa M, Engel T, Sollner TH, Rothman JE (2004) Countercurrent distribution of two distinct SNARE complexes mediating transport within the Golgi stack. Molecular biology of the cell 15 (4):1506–1518

70. Tai G, Lu L, Wang TL, Tang BL, Goud B, Johannes L, Hong W (2004) Participation of the syntaxin 5/Ykt6/GS28/GS15 SNARE complex in transport from the early/recycling endosome to the trans-Golgi network. Molecular biology of the cell 15 (9):4011–4022

71. D’Souza Z, Pokrovskaya I, Lupashin VV (2023) Syntaxin-5’s flexibility in SNARE pairing supports Golgi functions. Traffic 24 (8):355–379

72. Donaldson JG, Jackson CL (2011) ARF family G proteins and their regulators: roles in membrane transport, development and disease. Nature Reviews Molecular Cell Biology 12 (6):362–375

73. Schindler C, Chen Y, Pu J, Guo X, Bonifacino JS (2015) EARP is a multisubunit tethering complex involved in endocytic recycling. Nature cell biology 17 (5):639–650

74. Donaldson JG, Honda A, Weigert R (2005) Multiple activities for Arf1 at the Golgi complex. Biochimica et Biophysica Acta (BBA)-Molecular Cell Research 1744 (3):364–373

75. Fröhlich F, Petit C, Kory N, Christiano R, Hannibal-Bach H-K, Graham M, Liu X, Ejsing CS, Farese Jr RV, Walther TC (2015) The GARP complex is required for cellular sphingolipid homeostasis. Elife 4:e08712

76. Capasso S, Sticco L, Rizzo R, Pirozzi M, Russo D, Dathan NA, Campelo F, van Galen J, Hölttä-Vuori M, Turacchio G (2017) Sphingolipid metabolic flow controls phosphoinositide turnover at the trans-Golgi network. The EMBO journal 36 (12):1736–1754

77. Meissner JM, Bhatt JM, Lee E, Styers ML, Ivanova AA, Kahn RA, Sztul E (2018) The ARF guanine nucleotide exchange factor GBF1 is targeted to Golgi membranes through a PIP-binding domain. Journal of Cell Science 131 (3):jcs210245

78. Dmitriev OY, Patry J (2024) Structure and mechanism of the human copper transporting ATPases: Fitting the pieces into a moving puzzle. Biochimica et Biophysica Acta (BBA)-Biomembranes:184306

79. Lutsenko S (2010) Human copper homeostasis: a network of interconnected pathways. Current opinion in chemical biology 14 (2):211–217

80. Sluysmans S, Méan I, Xiao T, Boukhatemi A, Ferreira F, Jond L, Mutero A, Chang CJ, Citi S (2021) PLEKHA5, PLEKHA6, and PLEKHA7 bind to PDZD11 to target the Menkes ATPase ATP7A to the cell periphery and regulate copper homeostasis. Molecular biology of the cell 32 (21):ar34

81. Kaler SG (2011) ATP7A-related copper transport diseases—emerging concepts and future trends. Nature reviews Neurology 7 (1):15–29

82. Gale J, Aizenman E (2024) The physiological and pathophysiological roles of copper in the nervous system. European Journal of Neuroscience.

83. Hartwig C, Méndez GM, Bhattacharjee S, Vrailas-Mortimer AD, Zlatic SA, Freeman AA, Gokhale A, Concilli M, Werner E, Savas CS (2021) Golgi-dependent copper homeostasis sustains synaptic development and mitochondrial content. Journal of Neuroscience 41 (2):215–233

84. Comstra HS, McArthy J, Rudin-Rush S, Hartwig C, Gokhale A, Zlatic SA, Blackburn JB, Werner E, Petris M, D’Souza P (2017) The interactome of the copper transporter ATP7A belongs to a network of neurodevelopmental and neurodegeneration factors. Elife 6:e24722

85. Yi L, Kaler SG (2015) Direct interactions of adaptor protein complexes 1 and 2 with the copper transporter ATP7A mediate its anterograde and retrograde trafficking. Human molecular genetics 24 (9):2411–2425

86. Robinson MS, Antrobus R, Sanger A, Davies AK, Gershlick DC (2024) The role of the AP-1 adaptor complex in outgoing and incoming membrane traffic. Journal of Cell Biology 223 (7):e202310071

87. Kang H, Lee CJ (2024) Transmembrane proteins with unknown function (TMEMs) as ion channels: electrophysiological properties, structure, and pathophysiological roles. Experimental & Molecular Medicine 56 (4):850–860

88. Shin JJ, Crook OM, Borgeaud AC, Cattin-Ortolá J, Peak-Chew SY, Breckels LM, Gillingham AK, Chadwick J, Lilley KS, Munro S (2020) Spatial proteomics defines the content of trafficking vesicles captured by golgin tethers. Nature communications 11 (1):5987

89. Koreishi M, Gniadek TJ, Yu S, Masuda J, Honjo Y, Satoh A (2013) The golgin tether giantin regulates the secretory pathway by controlling stack organization within Golgi apparatus. PLoS One 8 (3):e59821

90. Casler JC, Papanikou E, Barrero JJ, Glick BS (2019) Maturation-driven transport and AP-1–dependent recycling of a secretory cargo in the Golgi. Journal of Cell Biology 218 (5):1582–1601

91. Laufman O, Freeze HH, Hong W, Lev S (2013) Deficiency of the Cog8 subunit in normal and CDG-derived cells impairs the assembly of the COG and Golgi SNARE complexes. Traffic 14 (10):1065–1077

