## Supplemental Figures 1-8 for "Acute GARP depletion disrupts vesicle transport, leading to severe defects in sorting, secretion, and O-glycosylation"

Figure S1 VPS54-myc-mAID expression rescues GARP-KO defects in HeLa cells

**A**

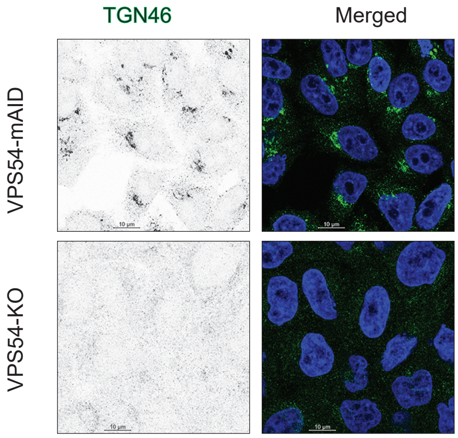


**B**

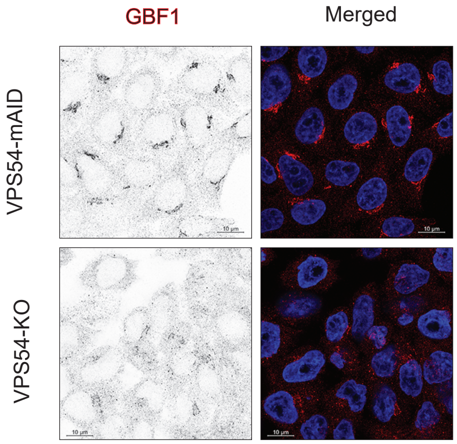


**C**

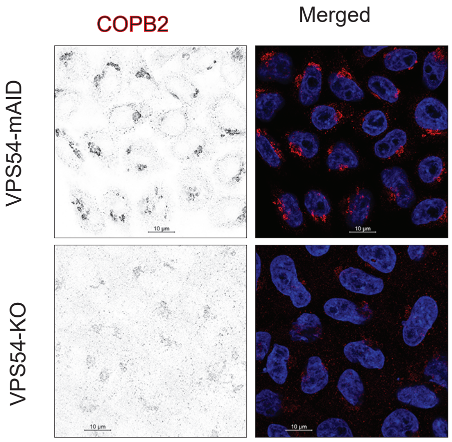

Figure S2 Rapid depletion of VPS54 in HeLa cells

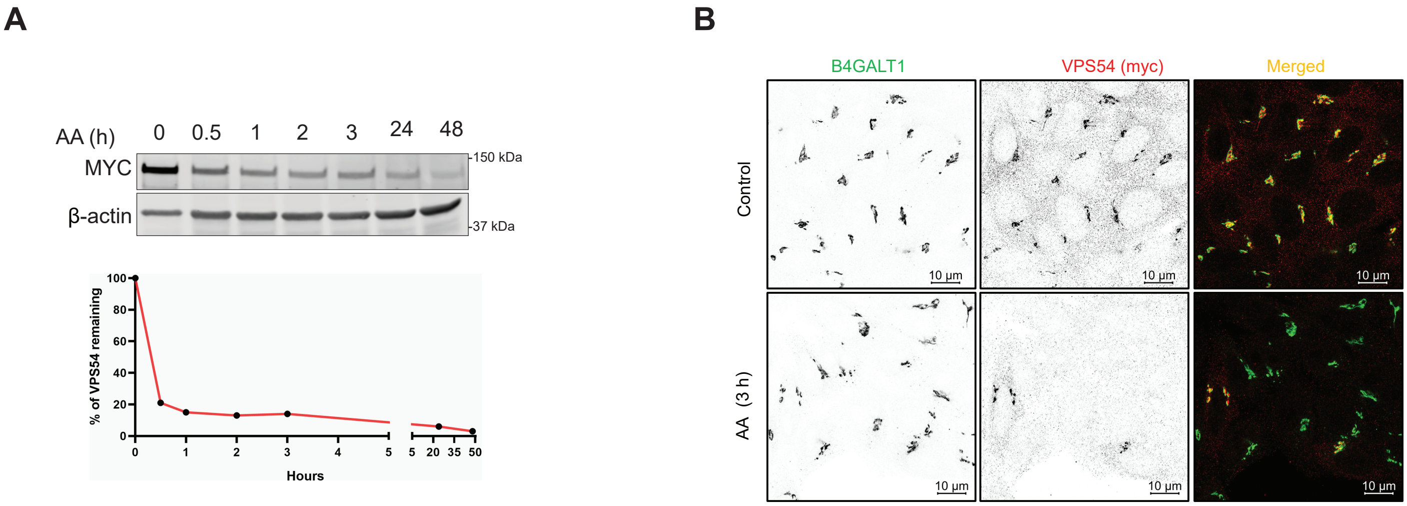

Figure S3 Effect of VPS54-KO on ATP7A and CPD localization

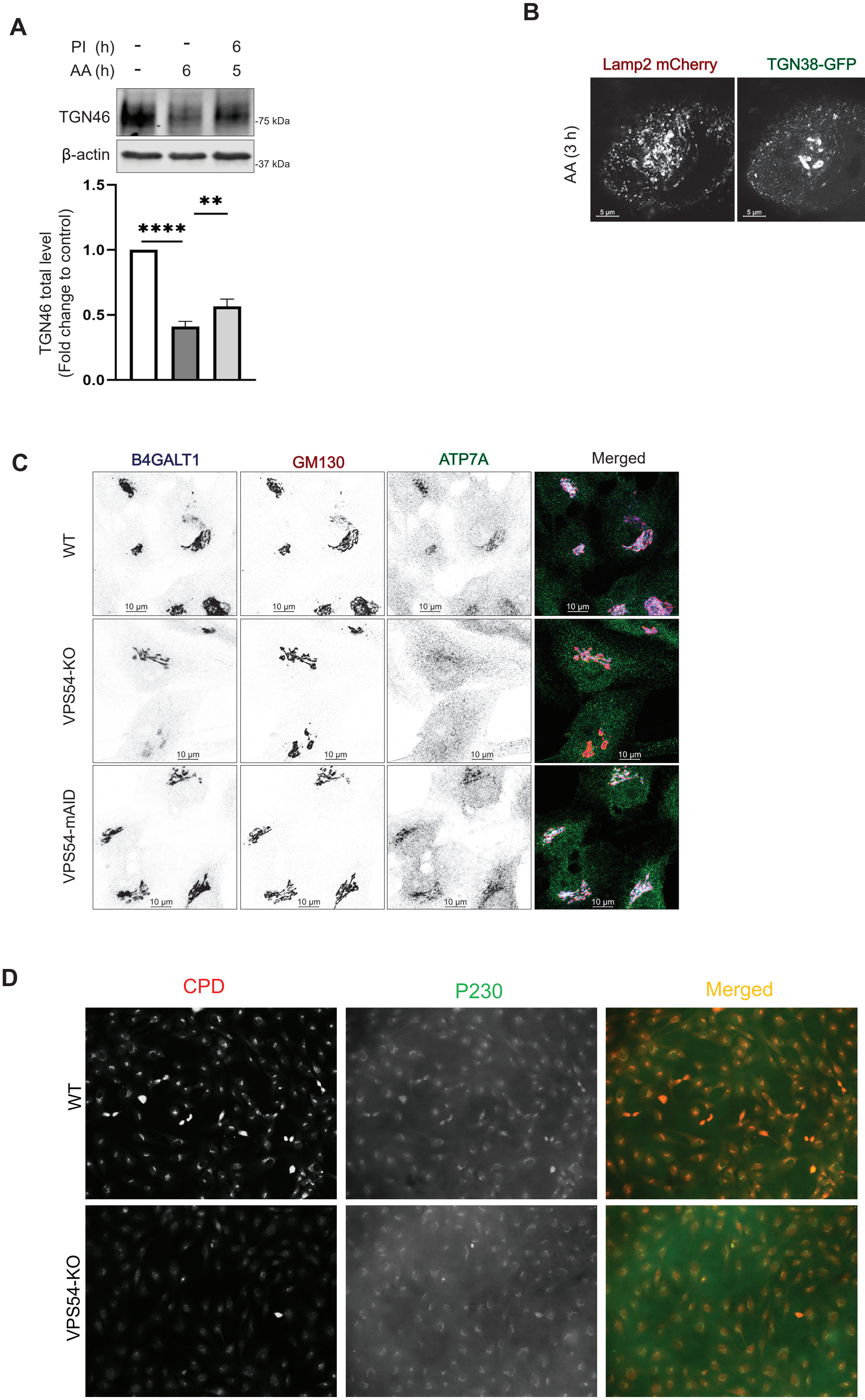

Figure S5  Acute VPS54 depletion do not affect MGAT1 and GALNT2

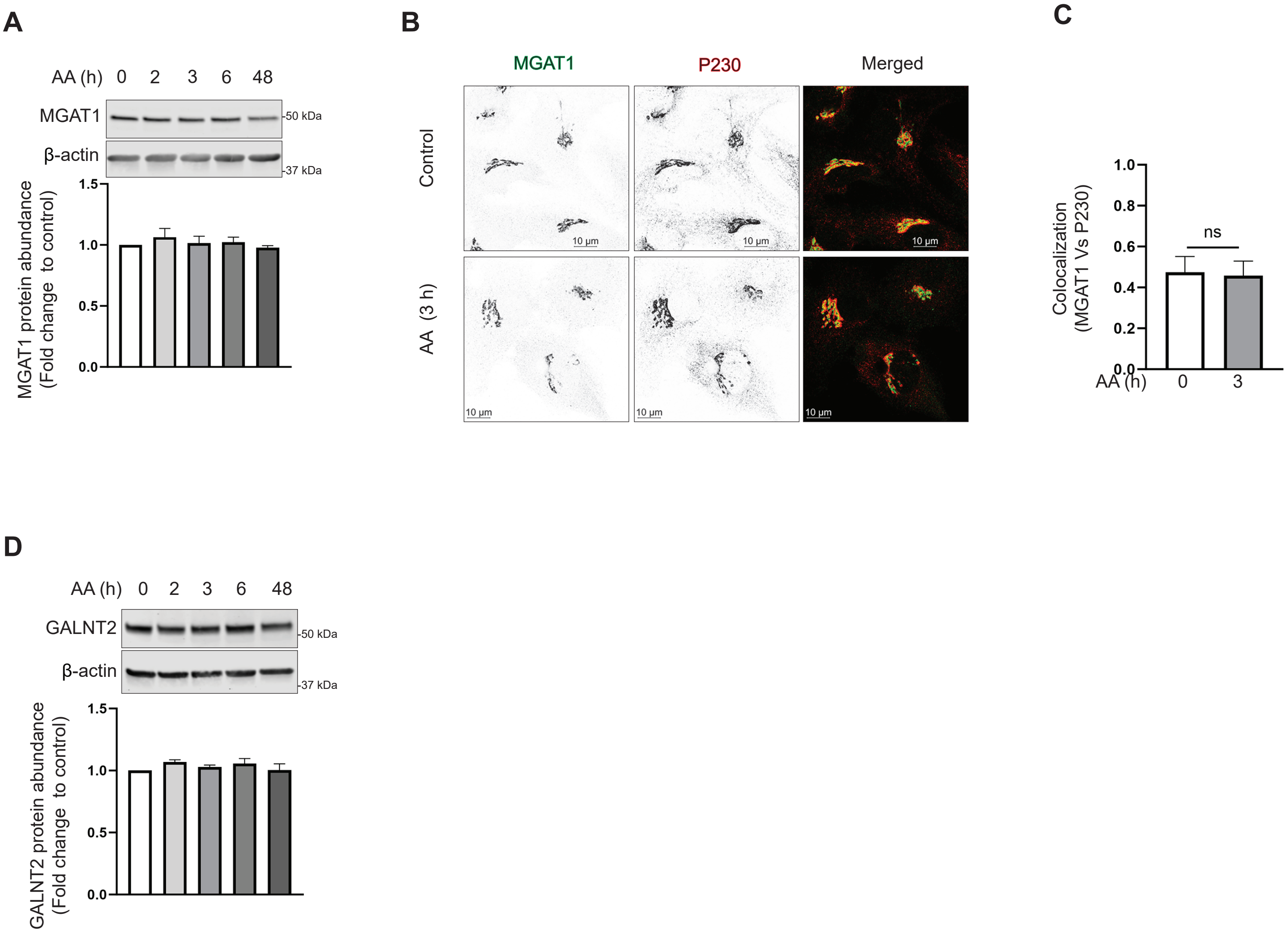

Figure S6 Rapid depletion of VPS54 stimulates B4GALT1 relocation to endosomes following CQ treatment

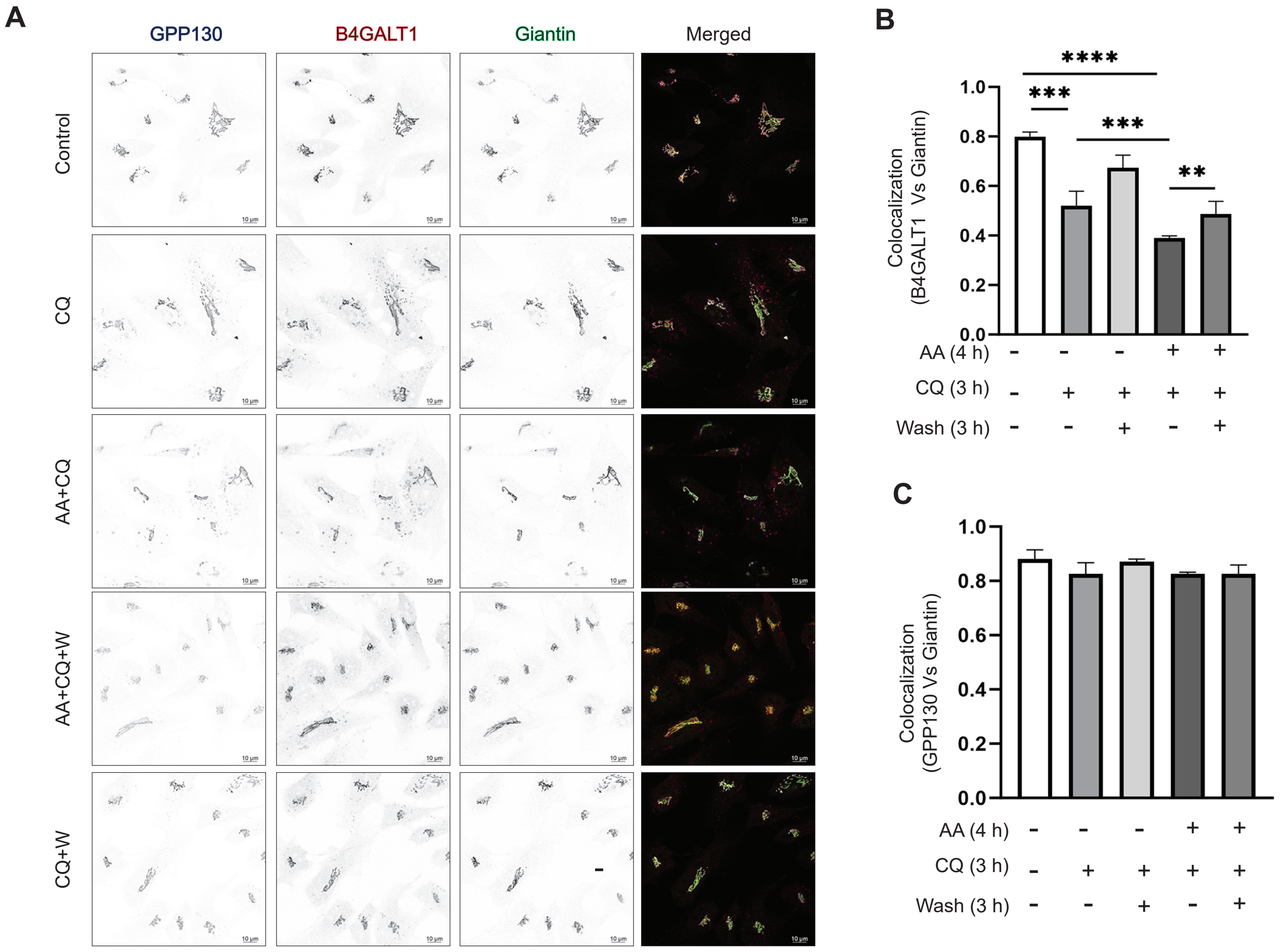

Figure S7 Acute VPS54 depletion do not affect BIG1 localization

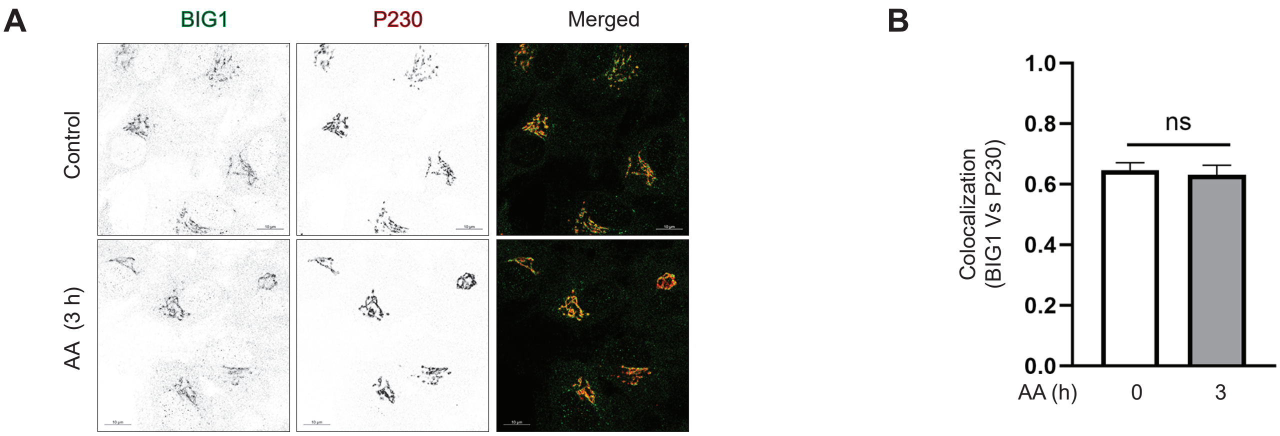

Figure S8 Increased GS15 vesicles after VPS54 degradation showed relocalization of C1GALT1 and STX6

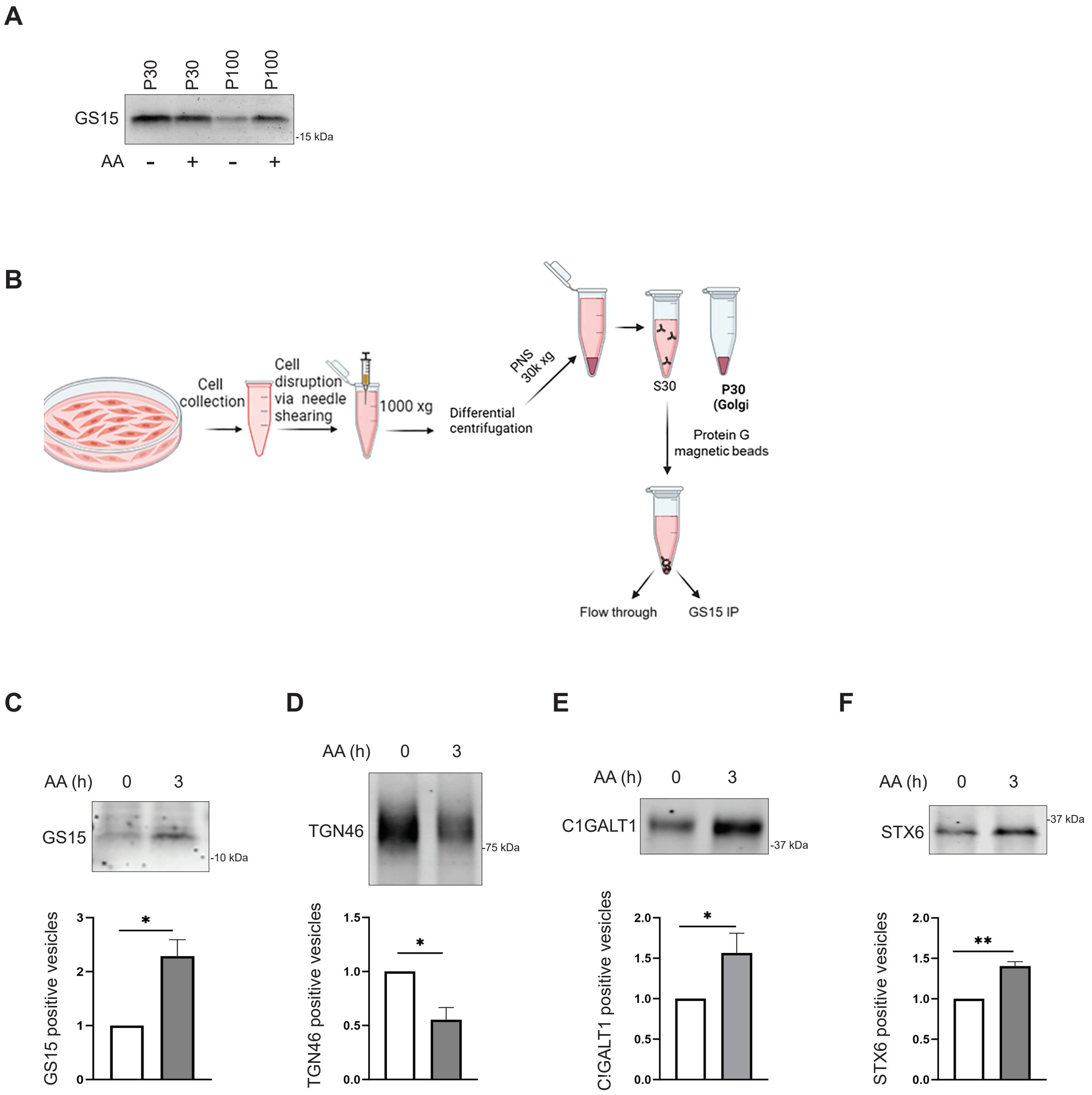
